# Gene regulation by DNA methylation is contingent on chromatin accessibility during transgenerational plasticity in the purple sea urchin

**DOI:** 10.1101/2021.09.23.461091

**Authors:** Samuel N Bogan, Marie E Strader, Gretchen E Hofmann

## Abstract

Epigenetic processes are proposed to contribute to phenotypic plasticity. In invertebrates, DNA methylation commonly varies across environments and can correlate or causally associate with phenotype, but its role in transcriptional responses to the environment remains unclear. Maternal environments experienced by the sea urchin *Strongylocentrotus purpuratus* induce 3 – 6x greater differential CpG methylation in offspring larvae relative to larval developmental environments, suggesting a role for DNA methylation in transgenerational plasticity (TGP). However, a negligible association has been observed between differentially methylated and differentially expressed genes. What gene regulatory roles does invertebrate DNA methylation possess under environmental change, if any? We quantified DNA methylation and gene expression in *S. purpuratus* larvae exposed to different ecologically relevant conditions during gametogenesis (maternal conditioning) or embryogenesis (developmental conditioning). We modeled differential gene expression and differential splicing under maternal conditioning as functions of DNA methylation, incorporating variables for genomic feature and chromatin accessibility. We detected significant interactions between differential methylation, chromatin accessibility, and genic architecture associated with differential expression and splicing. Observed transcriptional responses to maternal conditioning were also 4 – 13x more likely when accounting for interactions between methylation and chromatin accessibility. Our results provide evidence that DNA methylation possesses multiple functional roles during TGP in *S. purpuratus,* but its effects are contingent upon other genomic and epigenomic states. Singularly unpredictive of transcription, DNA methylation is likely one cog in the epigenomic machinery contributing to environmental responses and phenotypic plasticity in *S. purpuratus* and other invertebrates.

## 1. Introduction

Phenotypic plasticity, variation of a trait’s expression in the absence of genetic variation, can elicit both adaptive and maladaptive responses to rapid environmental change on ecological timescales (Donelan et al., 2020; Marshall & Uller, 2007). However, the molecular processes that mediate plasticity remain poorly understood. Uncovering these mechanisms will inform our understanding of physiological and evolutionary responses to changing environments (Herman & Sultan, 2011; Jones & Robinson, 2018). Epigenetic modifications to the genome such as DNA methylation are one suite of regulatory factors that, in some cases, underpin plasticity by driving changes in transcription and subsequent phenotypes (Xu et al., 2019). Our understanding of differential DNA methylation’s effects on gene expression and phenotype in metazoans is mostly derived from studies in vertebrates, but research in invertebrates finds correlations between changes in DNA methylation and phenotype across distinct environments, suggesting that methylation may influence acclimatization (Eirin-Lopez & Putnam, 2019; Hofmann, 2017). Connections between DNA methylation, plasticity, and acclimatization hinge on how and whether DNA methylation modulates gene regulation. Invertebrate DNA methylation frequently exhibits negligible relationships with differential expression (DE) and other modes of gene regulation in environmental studies, posing an obstacle to assessing the epigenetic basis of acclimatization to current and future environments. However, DNA methylation does not influence gene expression independent of other epigenetic and genetic factors. Using the purple sea urchin *Strongylocentrotus purpuratus* as a model invertebrate, we tested the hypothesis that the effects of differential methylation (DM) on gene regulation (differential expression and alternative splicing) are contingent upon additional epigenomic and genomic states such as chromatin accessibility and genic architecture. This integrated epigenomic approach allowed us to determine what regulatory roles DNA methylation in *S. purpuratus* possesses, if any, during plastic responses to ecologically relevant stressors that are worsening under climate change.

Multiple lines of evidence support DNA methylation’s influence on invertebrate ecology and biology. In broadly dispersing marine invertebrates, high connectivity reduces structure across populations inhabiting distinct environments, and interpopulation epigenetic divergence can exceed genetic divergence (Ardura, Zaiko, Moran, Planes, & Garcia-Vazquez, 2017; Johnson & Kelly, 2020; Ni et al., 2018; Watson, Baldanzi, Pérez-Figueroa, Gouws, & Porri, 2018; Zhang, Li, Kong, & Yu, 2018). *In situ* temporospatial environmental variation has been linked to modifications in invertebrate methylomes independent of genetic variation, demonstrating a potential role for DM during acclimatization. (Clark et al., 2018; Dimond & Roberts, 2020; Rodríguez-Casariego et al., 2020; Wang, Li, et al., 2021). DM of genes, gene modules, or whole genomes induced by environmental variation are associated with performance traits in stony corals, molluscs, crustaceans, and insects (Arsenault, Hunt, & Rehan, 2018; Clark et al., 2018; Li et al., 2018; Norouzitallab et al., 2014; Putnam, Davidson, & Gates, 2016). Causative tests of DNA methylation’s effect on phenotype have also been conducted. In the crustacean *Daphnia manga*, inhibition of *de novo* methyltransferases in the P generation induced genome-wide hypomethylation and DE of essential metabolic pathways, proceeding to impact performance and expression in F1 and F2 generations while leaving histone modifications unaffected (Lindeman et al., 2019; Vandegehuchte, Lemiere, Vanhaecke, Vanden Berghe, & Janssen, 2010). Environmentally induced changes and intraspecific variation in DNA methylation can also be inherited across the germline in some invertebrate taxa (Liew et al., 2020; Wang, Werren, & Clark, 2016). Testing the gene regulatory roles of DNA methylation will uncover its potential functions in a diversity of biological processes, particularly mechanisms of developmental and transgenerational plasticity (TGP) which drive acclimatization.

DNA methylation’s effects on gene regulation and its interactions with other epigenetic factors are both highly multiplicative. Most invertebrate phyla exhibit sparsely methylated genomes punctuated by high levels of CpG methylation within gene bodies (Keller, Han, & Yi, 2016; Suzuki, Kerr, De Sousa, & Bird, 2007; Zemach, McDaniel, Silva, & Zilberman, 2010). Patterns of invertebrate gene body methylation (GBM) vary across (i) genic features such as promoters, introns, exons, and UTRs (Li et al., 2018; Riviere et al., 2017) and (ii) phylogeny (de Mendoza et al., 2019; Keller et al., 2016; Sarda, Zeng, Hunt, & Yi, 2012). GBM positively correlates with gene expression in cnidarians (Dixon, Liao, Bay, & Matz, 2018; Li et al., 2018; Zemach et al., 2010), bivalve molluscs of *Crassostrea sp*. (Downey-Wall et al., 2020; Johnson, Sirovy, Casas, La Peyre, & Kelly, 2020), and arthropods (Bonasio et al., 2012; Gatzmann et al., 2018; Glastad, Gokhale, Liebig, & Goodisman, 2016; Kvist et al., 2018) with exceptions to this pattern evident in some species and cell types (de Mendoza et al., 2019; Flores et al., 2012; Wang, Song, et al., 2021). By contrast, inducible changes in invertebrate gene expression in response to environmental variation have frequently possessed insignificant relationships with differential GBM (Arsenault et al., 2018; Dixon et al., 2018; Downey-Wall et al., 2020; Johnson et al., 2020; Strader et al., 2020). In some arthropods, molluscs, and nematodes, there is evidence that GBM aids in regulating alternative splicing and exon skipping (Flores et al., 2012; Gao et al., 2012; Li-Byarlay et al., 2013; Libbrecht, Oxley, Keller, & Kronauer, 2016; Song, Li, & Zhang, 2017). Among these taxa, changes in alternative splicing under environmental change have shown relationships with differential GBM of varying strengths (Arsenault et al., 2018; Glastad et al., 2016). Invertebrate DNA methylation is also associated with chromatin state (Gatzmann et al., 2018; Nanty et al., 2011) and the suppression of spurious intragenic transcription (Li et al., 2018). Determining the function of DM during transcriptional responses to the environment thus requires an integrated approach that considers genic architecture, additional epigenetic features, and multiple modes of gene regulation (Moler et al., 2018).

The purple sea urchin *Strongylocentrotus purpuratus* is a uniquely poised model invertebrate in which to conduct an integrative test of DNA methylation’s regulatory roles during phenotypic plasticity. *S. purpuratus* is an abundant herbivore distributed throughout North America’s Pacific subtidal kelp forests and rocky intertidal. Populations inhabiting environmental gradients or mosaics exhibit genetic evidence of local adaptation and interpopulation variation in performance and gene expression under ecologically relevant stress (Evans, Chan, Menge, & Hofmann, 2013; Evans, Pespeni, Hofmann, Palumbi, & Sanford, 2017; Kelly, Padilla-Gamino, & Hofmann, 2013; Pespeni, Chan, Menge, & Palumbi, 2013; Pespeni & Palumbi, 2013). CpG methylation is more abundant in *S. purpuratus* relative to most invertebrates, likely because of its phylogenetic position as a basal deuterostome (Regev, Lamb, & Jablonka, 1998). TGP linked to maternal effects have been observed in *S. purpuratus* for traits including egg protein content, larval body size, gene expression, and DNA methylation (Hoshijima & Hofmann, 2019; Strader et al., 2020; Strader, Wong, Kozal, Leach, & Hofmann, 2019; Wong, Johnson, Kelly, & Hofmann, 2018; Wong, Kozal, Leach, Hoshijima, & Hofmann, 2019) alongside similar observations in congeneric *Strongylocentrotus spp.* (Ding et al., 2019) and other urchin genera (Clark et al., 2019; Karelitz, Lamare, Patel, Gemmell, & Uthicke, 2019; Wong & Hofmann, 2020; Wong & Hofmann, 2021). Maternal conditioning of *S. purpuratus* to abiotic conditions mimicking coastal upwelling can induce 3 – 6x greater DM in offspring larvae relative to the effects of larval development under upwelling (Strader et al., 2020; Strader et al., 2019). Maternal conditioning of *S. purpuratus* can also trigger DE of a larger number of genes than progeny conditioning (Wong et al., 2018). These results suggest a function for DM in facilitating TGP’s effects on gene expression, but negligible overlap between DM CpGs and DE genes has left that role ambiguous (Strader et al., 2020). Accounting for interactions between DNA methylation and additional epigenomic states in *S. purpuratus*, which may better explain epigenetic effects on transcription, is made possible by the species’ use as a model of deuterostome embryology, yielding developmental time series of chromatin accessibility (e.g., ATAC-seq) spanning the two-cell embryo to late prism larva.

To elucidate the gene regulatory roles of differential methylation during TGP, we quantified changes in DNA methylation, gene expression, and alternative splicing in prism larvae induced by maternal exposure to ecologically relevant, abiotic stress in *S. purpuratus* using data from Strader *et al.,* 2020 initially exhibiting limited overlap between DM and DE genes and robust annotations of chromatin accessibility during the *S. purpuratus* prism stage (Kudtarkar & Cameron, 2017). We then modeled differential expression and splicing as functions of DM, genic feature type, and chromatin accessibility to test the hypothesis that invertebrate DNA methylation’s regulatory role is contingent upon additional genomic and epigenomic factors and reveal epigenetic interactions influencing gene expression during TGP.

## 2. Methods

### 2.1. Data sources

To investigate the potential gene regulatory role of DNA methylation on transcription, we used previously published and publicly available datasets. For RNA-seq and bisulfite sequencing datasets, a controlled transgenerational experiment was performed (Strader et al. 2020). Briefly, adult urchins were conditioned to two treatments, non-upwelling (631 ± 106 µatm *p*CO2 and 16.8 ± 0.2 °C) and upwelling (1390 ± 307 µatm *p*CO2 and 12.7 ± 0.5 °C), mimicking variation in their natural environment (Hoshijima & Hofmann, 2019). Temperature and *p*CO_2_ conditions were maintained by a flow-through CO_2_ system (Fangue et al., 2010) and described in detail by Strader et al. 2020. Briefly, treated seawater was evenly pumped from two reservoir tanks to conditioning tanks at a rate of 20 L/hr. Adult urchins were induced to spawn and fertilizations were performed in ambient seawater conditions. Embryos were reared in either the same conditions as their parents or the reciprocal condition in triplicate using a flowthrough system with seawater treated as described above and by Strader et al. 2020.

Once larval development progressed to the early prism stage, replicate samples of 6,000 larvae were collected for RNA-seq and reduce representation bisulfite sequencing (RRBS) and flash frozen in liquid nitrogen before storage at -80 °C. Libraries for polyA-enriched RNA-seq and RRBS were constructed at the UC Davis genome center and sequenced on the Illumina 4000 (BioProject: PRJNA548926). The use of polyA-enriched RNA-seq libraries is beneficial for analyzing alternative splicing as it mitigates the contribution of unprocessed RNA to quantification of differential exon use (Sultan et al., 2014). RRBS poses fewer biases on CpG representation across genomic feature type relative to other reduced representation BS-seq methods (Trigg et al., 2021). ATAC-seq data was obtained through the GEO expression omnibus (BioProject: PRJNA377768). This dataset represents a developmental time course of Tn5 transposon chromatin accessibility regions in the *S. purpuratus* genome.

For comparison with the Strader *et al*., 2020 datasets, we chose ATAC-seq profiles for animals at 39 hours post-fertilization, the closest developmental time point for early prism larvae, for which 3 pooled samples were sequenced (GSM2520650, GSM2520651, GSM2520652). ATAC-seq bed files were concatenated and summarized using the R package *ChIPSeeker* v1.22.1 to quantify chromatin accessibility, expressed as the mean density of Tn5 ATAC-seq reads, across intra- and intergenic and genomic features. Mean chromatin accessibility of ± 500 bp transcriptional start sites (TSS), introns, and exons were each calculated in both gene- and feature-wise manners for analysis.

### 2.2. Gene expression analyses

RNA-seq reads were trimmed of adaptor sequences and filtered for quality using *TrimGalore*. Cleaned reads were mapped to the *S. purpuratus* genome (v3.1) using *hisat2* (Kim, Langmead, & Salzberg, 2015). Gene and exon counts were compiled with *featureCounts* (Liao et al. 2014) and analyzed in *edgeR* v3.28.1 (Robinson, McCarthy, & Smyth, 2010) for analyses of DE and differential exon use (DEU), a measure of exon inclusion or exclusion attributable to skipping and splicing. Gene-level and exon-level read counts were filtered to retain genes with > 0.5 counts per million (CPM) across at least 75% of all samples.

In order to test for DE and DEU, gene- and exon-level counts were modeled as a function of maternal environment, developmental environment, and their interaction using the robust iteration of the *edgeR* glmQLfit function to fit negative binomial generalized linear models (GLMs). Robust negative binomial dispersion estimates were calculated using empirical Bayesian shrinkage with the *edgeR* function estimateGLMRobustDisp. Log_2_ foldchanges (logFC), F-statistic scores, and p-values for genewise DE between maternal and developmental treatments were estimated using the *edgeR* function glmQLFTest to account for uncertainty of tagwise dispersion estimates and improve type I error control. Significant DE was determined using FDR-adjusted p-values (alpha = 0.05). DEU was assessed by applying the edgeR function diffSpliceDGE to exon-level negative binomial GLMs which output exon use coefficients denoted as ΔlogFC (exon logFC – gene logFC) as well as likelihood coefficients and FDR-adjusted p-values (alpha = 0.05) for LRTs of gene- and exon-level DEU (McCarthy, Chen, & Smyth, 2012; Robinson et al., 2010).

Quantifying DEU attributable to alternative splicing and exon skipping required the removal of genes exhibiting patterns of exon use consistent with spurious intragenic transcription and alternative TSS. Genes that are spuriously transcribed or exhibit alternative TSS possess exons with progressively lower inclusion toward 5’ ends (Li et al., 2018). Filtering out such genes from exon-level read counts used in DEU analysis required the fitting linear models to exon-use data of each gene and removing genes with positive slopes and a y-intercept of DEU > -0.25. Without this filtering step, 56.0% of genes that exhibited significant DEU under maternal upwelling would likely have been attributed to alternative TSS or spurious transcription while such genes would have composed 64.9% of significant DEU under developmental upwelling. While this approach honed in on DEU attributed to splicing and exon skipping, it is likely to remove genes with few exons in which a 5’ exon removed during splicing. Plots of DEU trends demonstrative of spurious transcription or alternative TSS are available in the GitHub repository https://github.com/snbogan/Sp_RRBS_ATAC.

Enriched gene ontologies (GO) were identified among genes exhibiting DE or DEU with Mann–Whitney U-tests input with signed, -log p-values using rank-based gene ontology analysis with adaptive clustering (Wright, Aglyamova, Meyer, & Matz, 2015) parameterized with alpha value = 0.05 and minimum GO-term group size = 5 genes for gene-level enrichment. Alpha = 0.01 and min. Minimum GO-term group size = 25 genes for exon-level enrichment to account for a mean exon count of ∼5 per gene in the *S. purpuratus* genome.

### 2.3. DNA methylation analyses

RRBS sequences were trimmed and filtered with *TrimGalore* specifying the –rrbs option. Trimmed RRBS reads were mapped to the genome using *Bismark* (Krueger & Andrews, 2011), and methylation calls were determined using the bismark_methylation_extractor command using default settings. Coverage files were used for subsequent DM analysis using an adapted *edgeR* workflow for RRBS data (Chen, Pal, Visvader, & Smyth, 2017). To examine feature-specific responses by DNA methylation to environmental treatments, DM was estimated as the logFC of summed methylation scores across all CpGs within the -1 kb promoters, introns, and exons of a given gene. For each feature type, summed counts were filtered to include only genes represented by ≥10 reads across all samples. edgeR was selected for DM analysis to provide a statistical framework unified with estimations of DE and DEU. Functional enrichment of GO terms among differentially methylated genes was assessed using Mann–Whitney U-tests input with signed, -log p-values using rank-based Gene Ontology analysis (Wright et al., 2015).

### 2.4. Modeling gene regulation as a function of epigenomic variation

Using a Bayesian framework, Gaussian linear models were fitted in order to predict baseline gene expression (log_2_ counts per million or logCPM) and binomial generalized linear models were fitted to binary values for the presence of alternative transcripts (e.g., splicing) as functions of mean CpG methylation and mean chromatin accessibility of promoters, introns, exons, and interactions between these predictors. Linear models of DE were fitted to predict logFC as a function of DM in -1 kb promoters, introns, and exons, as well as logCPM, chromatin accessibility across genic features, and components of genic architecture such as the total length of genic feature types. Linear models of DEU included predictors for DM of the corresponding exon, DM of all exons and introns of the associated gene, exon number, logCPM, chromatin accessibility across genic features, and genic architecture.

All linear and generalized linear models were fitted using the R package *brms* v2.14.0, an R interface to the Stan programming language for specifying Bayesian models (Bürkner, 2017). All models were fitted with scaled Z-score transformations of continuous variables. Linear models of DE and DEU were fit with studentized model families to reduce prediction of artificially high or low outcome variables and were specified with weakly informative normal priors (mean = 0; SD = 0.5) for both slope (β) and intercept parameters. Z-score transformations were used in order to improve model convergence and compare posterior distributions of β parameters for predictors of different dependent variables such as DE and DEU. Weakly informative priors expressing a low probability of DNA methylation affecting gene regulation accounted for knowledge that DM associated with plasticity has exhibited negligible singular effects on gene regulation in most invertebrates. Posterior distributions were sampled using 4 chains at 20,000 iterations each, including 5,000 warmup iterations.

Model selection was performed by (i) applying Bayes factors using the bayesfactor_models function in *bayestestR* v0.9.0 (Makowski, Ben-Shachar, & Lüdecke, 2019) to compare the likelihoods of models fit with iterative combinations of predictor variables excluding >3^rd^ order interactions and (ii) comparing the selected model to two additional, alternative models using k-fold cross validation via *rstanarm* v2.21.1 (Goodrich, Gabry, Ali, & Brilleman, 2020): a model of the outcome predicted by differential methylation alone and the selected model without its highest-order interaction term. Bayes factors are less likely to select complex models (Gronau & Wagenmakers, 2019) and were applied to a large number of varyingly complex models before the selected model’s predictive strength was evaluated with k-fold cross validation. To account for variation in RRBS read coverage across the data, the selected model was then refit to include an error parameter for estimated methylation and differential methylation that equaled the inverse CpG coverage of each gene or feature in the dataset. RRBS CpG coverage per feature is described in Supplemental Results for reported models. Posterior predictive checks were used to evaluate selected model predictions according to observed data. Effect significance was tested using probability of direction, a Bayesian corollary of the p-value (Makowski, Ben-Shachar, Chen, & Ludecke, 2019); fixed effects were deemed significant if 95% of their posterior distribution fell above or below 0. Inclusion Bayes factors were employed to test for the explanatory power of an effect by estimating the likelihood of observed data when fitted with a parameter relative to a null model excluding it (Hinne, Gronau, van den Bergh, & Wagenmakers, 2020). Diagnostic plots, QC information, and predictions of selected models are available in Supplemental Results. The specifications and relative likelihoods of selected and unselected models are available in the following GitHub repository: https://github.com/snbogan/Sp_RRBS_ATAC.

## 3. Results

The results of our study demonstrate (i) that differential DNA methylation likely bears gene regulatory effects during TGP in *Strongylocentrotus purpuratus* and (ii) that these effects are conditional upon chromatin accessibility and genic architecture. We observed positive correlations between baseline DNA methylation, transcript abundance, and the presence of alternative splice forms within genes. With regard to plastic changes in DNA methylation and gene regulation, differential GBM interacted with chromatin accessibility and genic architecture to affect both differential gene expression and differential exon use/splicing such that the strength and direction of DM’s effects were contingent upon these additional genomic and epigenomic states. We describe these results in three sections below, focusing first on baseline relationships between DNA methylation and transcription, followed by epigenetic and gene regulatory responses to experimental upwelling. Finally, we present the results of integrated epigenomic models of DNA methylation’s gene regulatory effects during TGP.

### 3.1. Associations between constitutive epigenomic states and transcription

GBM in *S. purpuratus* prism larvae showed significant and positive correlations with gene expression level and the probability of associated alternative transcriptional variants. CpGs within -1 kb promoters, exons, and introns exhibited mean methylation levels of 34.70%, 43.21%, and 44.64%, respectively. Mean promoter methylation demonstrated a significant, albeit weak, negative effect on the expression (logCPM) of corresponding genes. By contrast, mean exon and intron methylation both exhibited stronger, positive effects on expression. Genes that were highly methylated in either exons or introns were ∼2x more expressed than unmethylated genes (Fig. 1A). Exon and intron methylation also exhibited a significant, antagonistic interaction such that genes with high methylation at both introns and exons were not more expressed than genes with high methylation at only introns or exons. Interestingly, accounting for TSS accessibility in models of logCPM resulted in the loss of a significant effect of intron methylation on gene expression. Lastly, genes with high levels of intron or exon methylation were more likely as to exhibit transcript variants consistent with alternative splicing, alternative TSS, and/or exon skipping. The relationship between the probability of transcript variants and methylation at promoters was insignificant (Fig. 1B).

**Figure 1.**
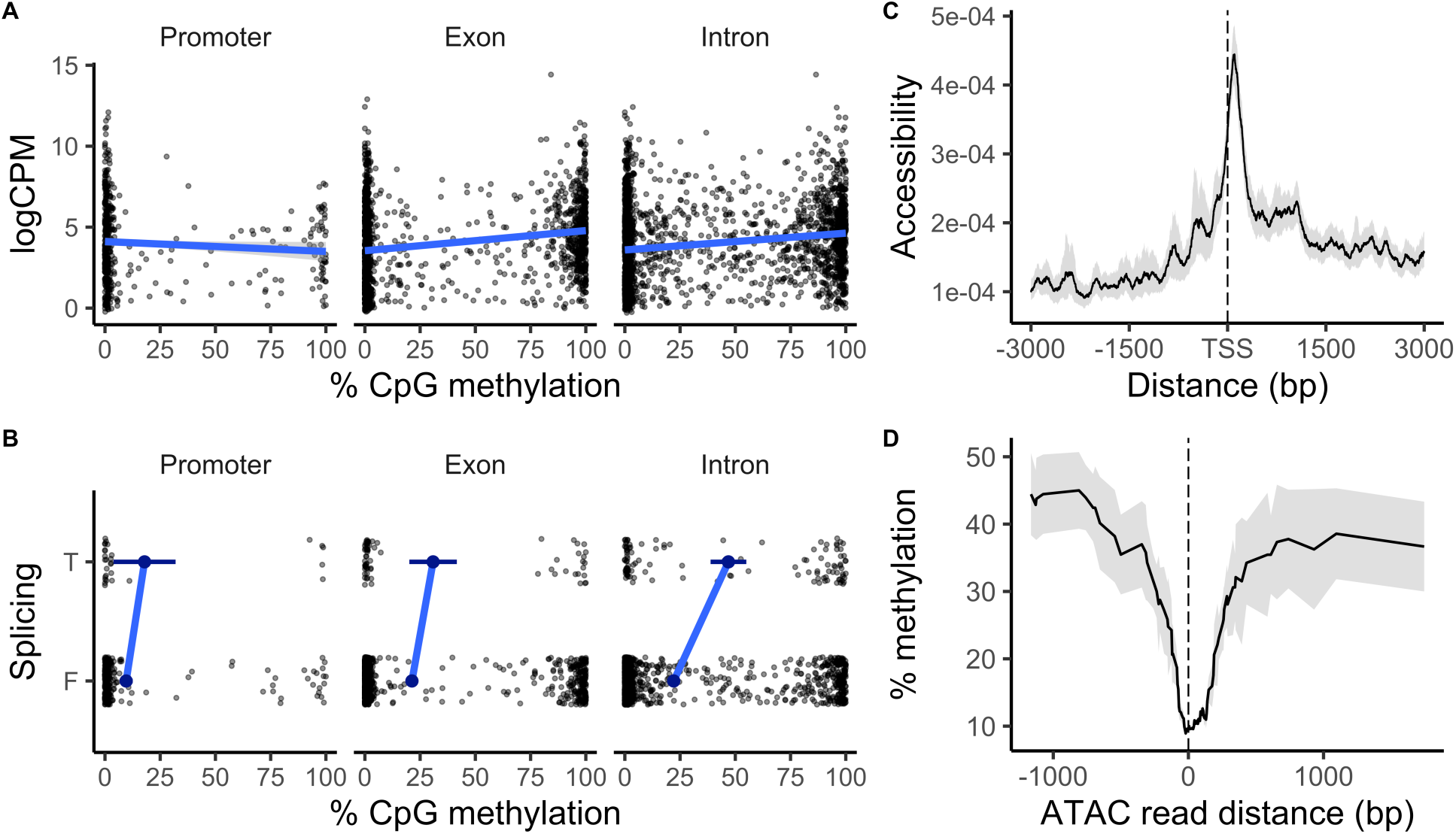
Relationships between constitutive DNA methylation, expression, and chromatin accessibility in larval *Strongylocentrotus purpuratus*. (a) Median methylation averaged across CpGs in the promoter, introns, and exons per gene are plotted against transcript abundance and (b) whether a gene exhibited transcript variants. Error bars depict ± 95% CI. (c) Loess trend of mean ATAC-seq read density (i.e., chromatin accessibility) ± 95% CI is plotted across distance to TSS sites. (d) 50 bp sliding window averages of CpG methylation ± 95% CI are plotted across distance to accessible chromatin regions for which ATAC-seq reads intersect from all replicate samples.

Chromatin accessibility at TSS, exons, and introns were also correlated with gene expression level, but not with the probability of transcriptional variants. However, selected models of logCPM did not include parameters related to chromatin accessibility and such parameters also yielded low inclusion Bayes factors. Chromatin accessibility was enriched proximal to TSS (Fig. 1C) but was greatest in introns, which exhibited a mean of 0.19 ± 0.11 ATAC-seq reads per bp compared to 0.043 ± 0.001 and 0.039 ± 0.043 in TSS and exons, respectively. Open chromatin regions showed ∼30% less CpG methylation than inaccessible regions (Fig. 1D). Gene-level intron and exon methylation showed no relationship with chromatin accessibility of either genic feature (Fig. S1A). Chromatin accessibility within ± 500 bp of TSS and gene bodies both showed significant and positive correlations with gene expression, with TSS accessibility exhibiting an effect that was 32.98% stronger than gene body accessibility (Fig. S1B). TSS accessibility did exhibit a significant and positive correlation with the probability of alternative transcriptional variants. This effect was insignificant after accounting for intron methylation however. Thus, intron methylation was the only significant predictor of alternative splicing events.

### 3.2. Transcriptional and epigenetic responses to environmental variation

Maternal and developmental exposure to experimental upwelling induced DE, as well as DEU consistent with alternative splicing and exon skipping in prism larvae of *S. purpuratus*. As reported by Strader et al. 2020, differential CpG methylation was observed in response to maternal upwelling, but no DM was observable under developmental upwelling. This distinction between maternal and developmental effects remained constant after quantifying DM averaged across genes’ promoters, exons, and introns.

Developmental upwelling exposure induced 2,263 upregulated and 2,459 downregulated, differentially expressed genes (DEGs). Maternal exposure induced 1,380 upregulated and 1,025 downregulated DEGs (Fig. 2). After applying a log_2_FC cutoff of > 1.0, 309 significant developmental DEGs were retained while 245 maternal DEGs were retained. Although the developmental treatment gave rise to a greater number of DEGs, absolute logFCs of DE among maternal DEGs were significantly higher than absolute logFCs of developmental DEGs by 10.45%. Functional enrichment of biological process, molecular function, cellular component GO terms among DEGs was observed in response to maternal and developmental treatments and is extensively reported by Strader et al. 2020.

**Figure 2.**
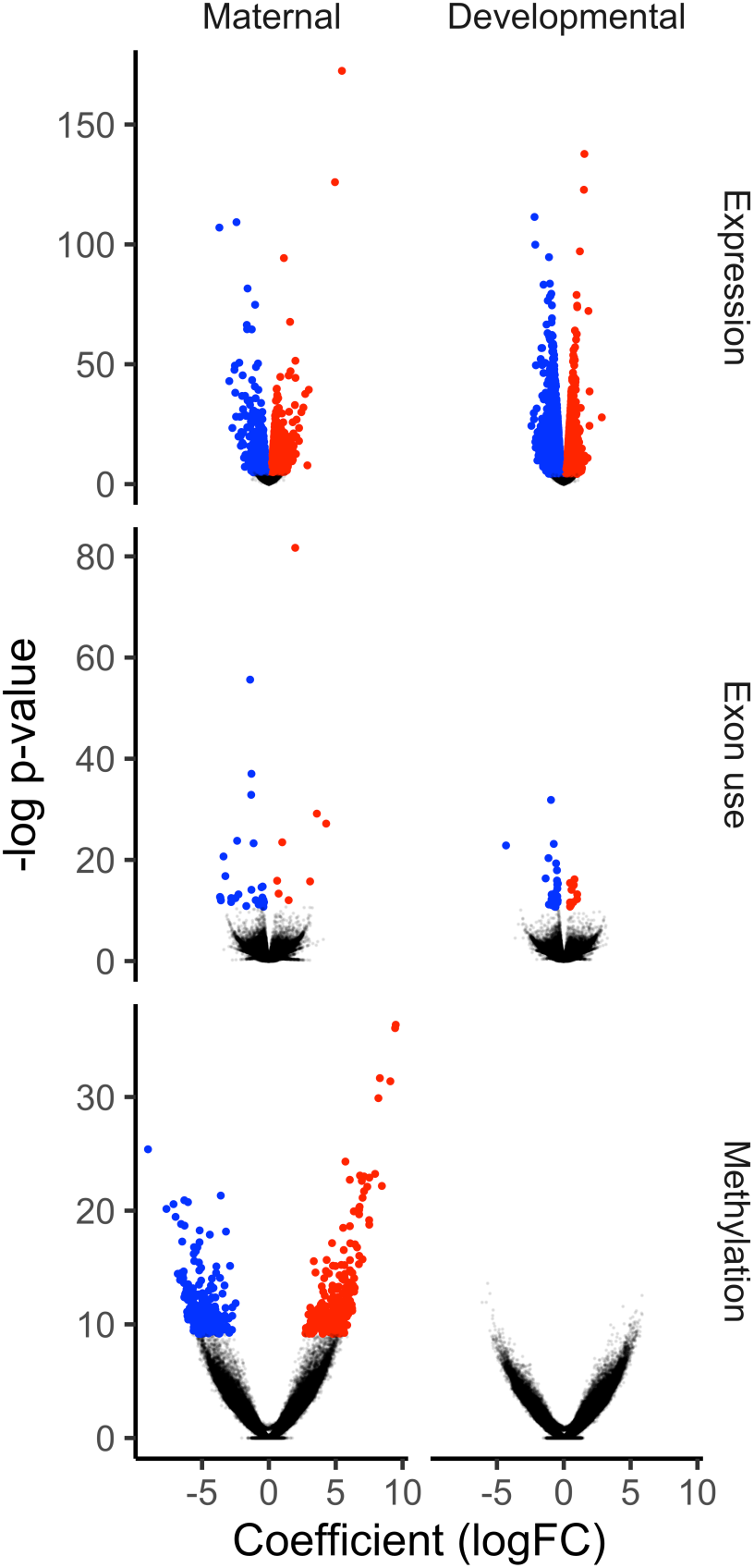
Molecular responses to developmental and maternal upwelling exposure. (top) Volcano plots of differential expression depicting genewise -log_2_ p-values, log_2_FC, and significant differential expression (color). (middle) Volcano plots of differential exon use or DEU (e.g., splicing) depicting exon-level -log_2_ p-values, DEU coefficients, and significant DEU (color). (bottom) Volcano plots of differential CpG methylation depicting CpG-level -log_2_ p-values, log_2_FC of differential methylation, and significant differential methylation (color). Red and blue points depict significant positive and negative coefficients, respectively (FDR < 0.05).

Substantially less DEU occurred in response to experimental upwelling relative to DE. Significant DEU was evaluated using both gene- and exon-level tests. Developmental upwelling induced 78 differentially spliced genes (DSGs) while maternal upwelling induced 121 DSGs, with 16 DSG genes shared between treatments. 43 and 49 genes were both differentially expressed and differentially spliced in response to developmental upwelling and maternal upwelling, respectively (Fig. S2). Significant DEU was detected among 44 exons in response to developmental upwelling: 14 upregulated or “included” exons and 30 downregulated or “dropped” exons. DEU in response to maternal upwelling occurred in 47 exons: 12 included and 35 dropped exons (Fig. 2). The molecular function (MF) GO terms ‘structural molecule activity’ and ‘structural constituent of ribosome’ and the biological process terms ‘obsolete GTP catabolic process’, ‘small GTPase mediated signal transduction’, and ‘cellular amide metabolic process’ were enriched among differentially spliced exons in response to both maternal and developmental treatments. Exons differentially spliced under the maternal treatment were also enriched for the biological processes (BP) ‘cellular localization’ and ‘nuclear transport’ among others (see Supplemental Material).

In response to maternal upwelling, 288 CpGs were hypermethylated and 233 were hypomethylated. Zero CpGs were differentially methylated in response to developmental upwelling (Fig. 2). This effect was distinct from DE and DEU, which both exhibited greater or equal variation in response to developmental upwelling compared to maternal upwelling. Functional enrichment among differentially methylated genes is described in detail by Strader et al 2020.

### 3.3. Chromatin state, genic architecture, and differential methylation interactively influenced transcriptional responses to environmental variation

The strength of differential GBM’s effect on DE was conditional upon chromatin accessibility and transcript abundance. Differential GBM across whole genes did not affect DE, while intron DM was significantly associated with DE. The selected model of DE as a function of intron DM under maternal upwelling included a significant three-way interaction between intron DM, TSS accessibility, and logCPM. Intron DM had higher absolute effects on DE among genes with poorly accessible TSS. These effects were positive among genes with low expression and negative for highly expressed genes (Fig. 3A). The inclusion Bayes factor for the interaction between intron DM, logCPM, and TSS accessibility was 3.84 (Fig. 3B). A Fishers exact test demonstrated that genes in the lowest logCPM quartile and lowest quartile of TSS accessibility were enriched with MF GO terms that included ‘nucleotidyltransferase activity’ and ‘cytoskeletal motor activity’, two MF terms that were also enriched among genes with CpGs that were differentially methylated under maternal upwelling (Strader et al., 2020). The biological process GO terms ‘plasma membrane bounded cell projection assembly’ and ‘movement of cell or subcellular component’ were also enriched among genes with low TSS accessibility and expression, among others (see Supplemental Material). Genes in the lowest TSS accessibility quartile and highest logCPM quartile were enriched with the MF terms ‘threonine-type endopeptidase activity’, ‘transcription regulator activity’, and ‘calcium ion binding’. An Intron DM standard deviation of +1.00 corresponded to a DE standard deviation of +0.26 ± 0.12 among genes in the lowest quartile of TSS accessibility and lowest logCPM quartile and -0.14 ± 0.09 for low TSS accessibility, high logCPM genes. Lastly, some model iterations in the top 25% of marginal likelihoods indicated that intron length interacted with intron DM to affect expression. Intron hypermethylation of genes in the top quartile of intron length silenced expression while genes in the middle-to-lowest quartiles of intron length showed enhanced expression as a result of intron hypermethylation (Fig. S3), though the inclusion Bayes factor for this interaction was negligible (BF = 1.67).

**Figure 3.**
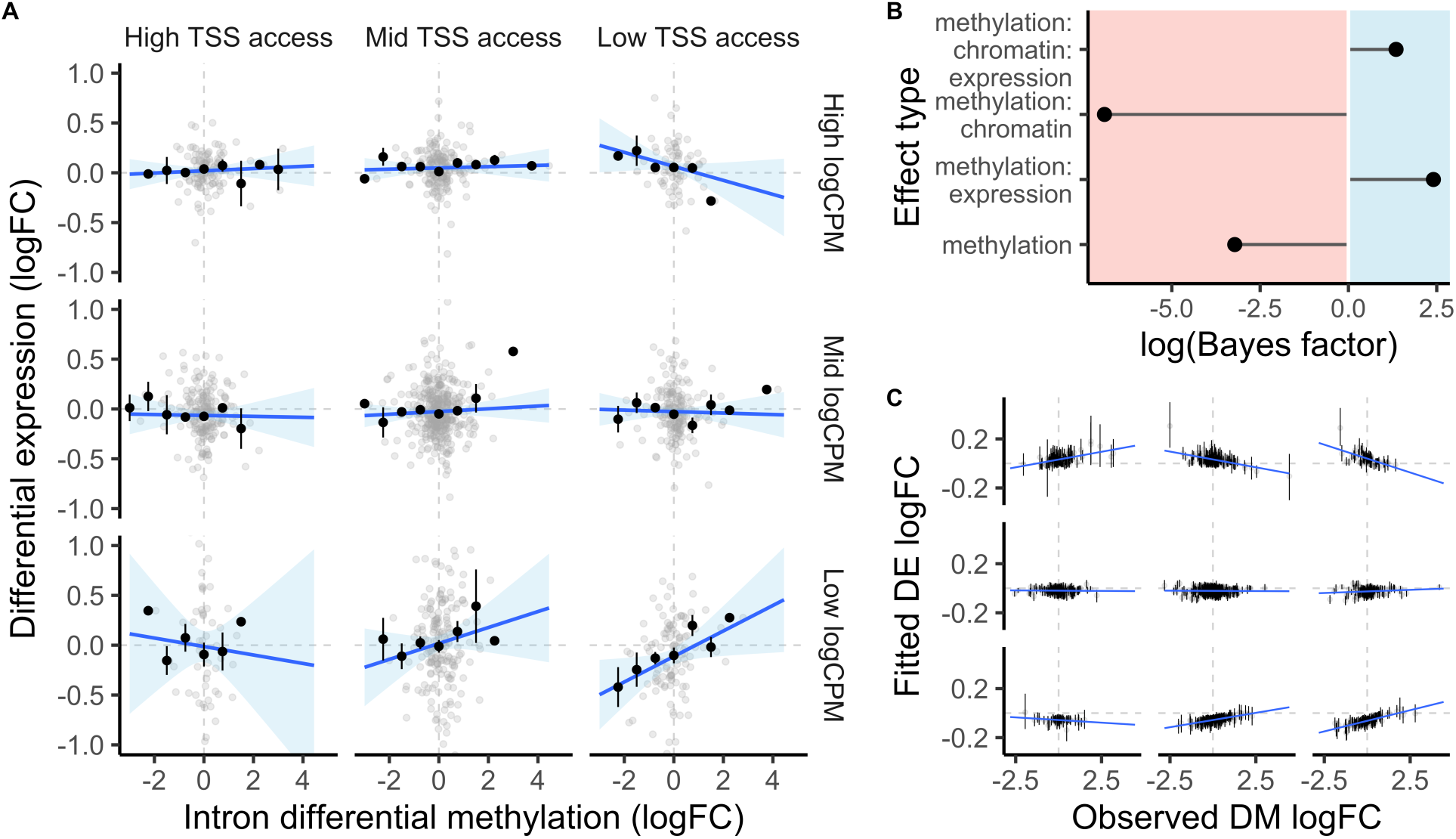
Differential intron methylation affected expression conditional upon TSS accessibility and transcript abundance. (a) Differential gene expression under maternal upwelling is plotted against mean intron differential methylation. Rows and columns group data based on transcript abundance and TSS accessibility quartiles, respectively. First and last rows/columns denote highest and lowest quartiles. Linear regressions are fitted across observed values. Average logFC across binned intron differential methylation is plotted as black points ± SE. (b) Log-scale inclusion Bayes factors depicting the probability of observed data under a parameter, including the interaction plotted in panel A. (c) logFC values fitted by the selected model of differential expression as a function of intron differential methylation. Error bars represent ± 95% credibility intervals. Columns, rows, and the x-axis depict binned TSS accessibility, logCPM, and observed intron differential methylation as presented in panel A.

Differential exon methylation responding to maternal upwelling interacted with gene body accessibility and genic architecture to affect DEU attributable to variation in alternative splicing and/or exon skipping. Selected models of DEU yielded a significant three-way interaction between exon DM, exon accessibility, and the total genic intron length. Both the strength and direction of exon DM’s effect on DEU was conditional upon exon accessibility and intron length. Positive correlations between exon DM and DEU were observed among exons from genes with poor exon accessibility while negative correlations were observed among exons from genes with highly accessible exons. Absolute effect strengths of exon DM on DEU were stronger among genes with longer introns (Fig. 4A). The inclusion Bayes factor of exon DM, exon accessibility, and intron length’s interactive effect on DEU equaled 13.03 (Fig. 4B). Exons from genes with long introns in the lowest quartile of exon accessibility exhibited a Z-score β of +0.43 ± 0.20. Exons from genes with long introns in the highest quartile of exon accessibility bore an effect strength of -0.12 ± 0.14. Genes in the lowest quartile of exon accessibility and highest intron length quartile were enriched with the MF GO terms ‘calcium ion binding’, ‘cytoskeletal motor activity’, and ‘ATPase activity’, among others, and the BP terms ‘purine-containing compound metabolic process’, ‘microtubule-based movement’, and ‘cell adhesion’. Genes with long introns and high exon accessibility were also enriched with the MF term ‘calcium ion binding’, as well as ‘transporter activity’, ‘small molecule binding’, and others, and enriched BP terms including ‘cell adhesion’, ‘localization’, ‘regulation of intracellular signal transduction’ (see Supplemental Material).

**Figure 4.**
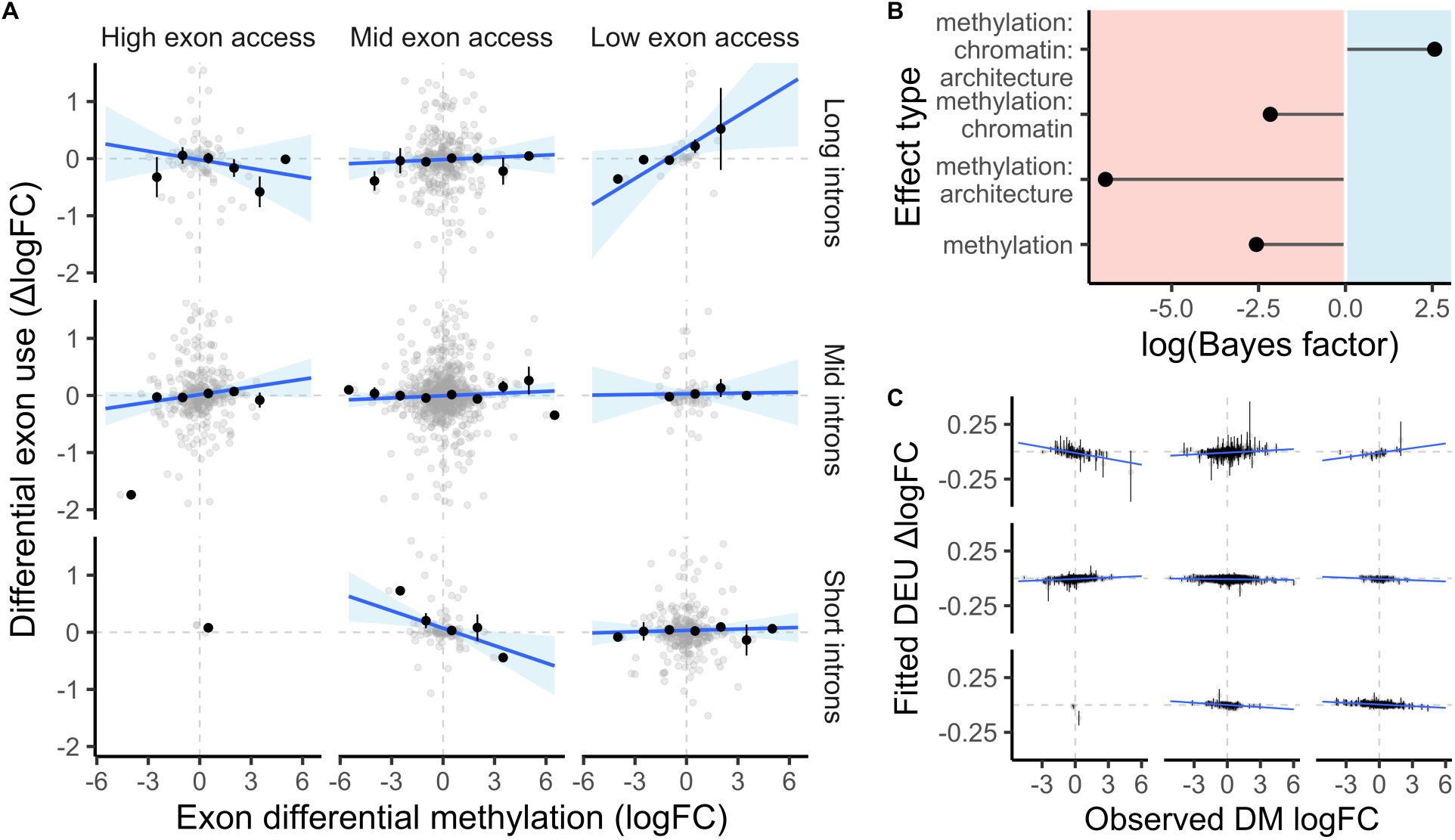
Differential exon methylation affected splicing conditional upon exon accessibility and genic architecture. (a) Differential exon use under maternal upwelling is plotted against mean exon differential methylation. Rows and columns group data based on total genic intron length and exon accessibility quintiles. First and last rows/columns denote highest and lowest quintiles. Linear regressions are fitted across observed values and are absent in the bottom left panel due to a low representation of observations. Average ΔlogFC across binned exon differential methylation is plotted as black points ± SE. (b) Log-scale inclusion Bayes factors depicting the probability of observed data under a parameter, including the interaction plotted in panel A. (c) ΔlogFC values fitted by the selected model of differential exon use as a function of exon differential methylation. Error bars represent ± 95% credibility intervals. Columns, rows, and the x-axis depict binned exon accessibility, intron length, and observed exon differential methylation as presented in panel A. Points represent exons in panels A and C.

## 4. Discussion

We sought to characterize relationships between differential methylation and transcriptional plasticity in the purple urchin *Strongylocentrotus purpuratus*, a species for which DNA methylation appears to play a role in transgenerational plasticity (Strader et al., 2020; Strader et al., 2019; Wong et al., 2019). *S. purpuratus* induces both significant differential expression and differential exon use (e.g., alternative splicing) in response to developmental and maternal exposure to experimental upwelling. Furthermore, differential gene body methylation in *S. purpuratus* larvae induced by maternal exposure to this ecologically relevant stressor exhibits significant and strong effects on DE and DEU among subsets of genes contingent upon chromatin accessibility and genic architecture (e.g., genomic feature type and genic feature length). Observed changes in expression and exon use under maternal conditioning were 4x and 13x more likely, respectively, when accounting for interactions between differential GBM, chromatin accessibility, and genic architecture. These results support the hypotheses that DM induced during TGP elicits multiple gene regulatory effects in *S. purpuratus* and, secondly, that these effects are conditional upon genomic and epigenomic features extrinsic of DNA methylation. Here we (i) discuss potential mechanisms to explain interactions between DNA methylation, chromatin accessibility, and genic architecture, (ii) interpret our results in the context of *S. purpuratus* and invertebrate physiological ecology, and (iii) highlight questions to be pursued in future studies of ecological epigenomics in metazoans.

### 4.1. Relationships between DNA methylation, chromatin accessibility, and gene expression in *Strongylocentrotus purpuratus* and other invertebrates

Baseline patterns of genomic methylation, chromatin accessibility, transcription, and the relationships between these processes in *S. purpuratus* were consistent with those typically observed in other invertebrate lineages (Bonasio et al., 2012; Dixon et al., 2018; Downey-Wall et al., 2020; Flores et al., 2012; Gao et al., 2012; Gatzmann et al., 2018; Glastad et al., 2016; Johnson et al., 2020; Kvist et al., 2018; Li et al., 2018; Li-Byarlay et al., 2013; Libbrecht et al., 2016; Song et al., 2017; Zemach et al., 2010). Our results also further illuminate the relationship between baseline intragenic methylation and gene expression in metazoans; we observed an antagonistic relationship between intron and exon methylation in *S. purpuratus* such that a saturation of methylation in one genic feature reduced the additive effect of methylation on expression by the other. Exon methylation was significantly correlated with gene expression regardless of intron methylation, but the effect of intron methylation was conditional upon low exon methylation. Several competing and non-competing hypotheses have been put forward to explain positive correlations between GBM and gene expression. Firstly, GBM inhibition in metazoans can reduce gene expression, causally linking GBM and expression (Lindeman et al., 2019; Yang et al., 2014). Recent advances in invertebrate epigenomics suggest that intragenic DNA methylation is bound by methyl-DNA-binding domain protein 2/3, which recruits acetyltransferases to promote H3K27 acetylation and initiate transcriptional elongation (Xu et al., 2021). Non-competing hypotheses posit that GBM may correlate with gene expression because it supports sequence conservation and transcriptional homeostasis. For example, genes with intragenic hypermethylation (i) are less accessible on average in at least some invertebrates (Gatzmann et al., 2018), which can protect them from mutation (Shi et al., 2016), and (ii) evolve at a slower rate across invertebrate lineages (Hunt, Brisson, Yi, & Goodisman, 2010; Park et al., 2011; Sarda et al., 2012). Intragenic methylation can also ensure a reduction in spurious transcription at noncanonical TSS, promoting transcriptional homeostasis (Neri et al., 2017). The singular effect of exon methylation on baseline expression in *S. purpuratus,* and the absence of a singular effect attributed to introns, suggest a unique association between exon methylation and expression. Conversely, the antagonistic interaction between exon and intron methylation may be indicative of functional redundancy for a secondary association between GBM and gene expression. For example, it is possible that exon methylation is more strongly associated with reduced intragenic accessibility, preventing mutation in coding regions of conserved genes, while exon and intron methylation may contribute relatively equally in directing transcriptional elongation to canonical TSS by preventing intragenic accessibility.

Intron methylation was also positively correlated with the prevalence of transcript variants among *S. purpuratus* genes consistent with associations between GBM and alternative splicing in other metazoans. Intragenic hypermethylation recruits methyl CpG binding protein 2 to splice junctions, which promotes exon recognition and, in some cases, intron retention via alterations to the elongation rate of RNA pol II (Maunakea, Chepelev, Cui, & Zhao, 2013; Wong et al., 2017). Considering existing evidence that exon methylation is positively associated with exon inclusion, it is surprising that intron methylation was associated with splicing in *S. purpuratus* while exon methylation was not. Relationships between the inclusion of specific exons or introns and their baseline methylation may still exist in *S. purpuratus*. We chose to model genewise averages of median CpG methylation at exons and introns as predictors of splice variation rather than fitting models of exon- or intron-specific use. This decision was made because our short read RNA-seq approach allowed for analyses of DEU between treatment groups, but not baseline exon use independent of treatment; we could not determine whether a methylated exon or intron was more likely to be constitutively skipped or included in a transcript. A more resolute quantification of GBM’s effect on alternative splicing in *S. purpuratus* or other invertebrates would be achieved by incorporating BS-seq with long read RNA-seq to enable isoform-specific read counting.

Patterns of chromatin accessibility and its relationship with gene expression in *S. purpuratus* were typical of other invertebrates and metazoans. Models of logCPM that included TSS or exon accessibility as predictors demonstrated significant, positive correlations with gene expression. However, variables related to chromatin accessibility were not included in the selected model of logCPM and yielded low Bayes factors for inclusion. Indeed, chromatin accessibility is necessary for the initiation of transcriptional elongation at canonical TSS (Klemm, Shipony, & Greenleaf, 2019). No relationships were observed between chromatin accessibility at genic features and the presence of transcript variants. Alternatively spliced exons exhibit more accessibility and less nucleosome occupancy than constitutively retained exons (Naftelberg, Schor, Ast, & Kornblihtt, 2015), inconsistent with the lack of relationship between chromatin accessibility and the prevalence of transcript variants in *S. purpuratus*. Finally, our observation that CpG methylation precipitously declined proximal to open regions of chromatin is consistent with findings in other invertebrates, vertebrates, and plants (Gatzmann et al., 2018; Lhoumaud et al., 2019; Zhong et al., 2021) and is characteristic of methylation CpG binding protein’s interactions with repressive chromatin factors (Klose & Bird, 2006).

### 4.2. Differential methylation’s effect on expression depends on chromatin state and genic architecture

Our finding that DM induced by maternal environment affected DE and DEU conditional upon chromatin accessibility and genic architecture supports the hypotheses (i) that DM possesses gene regulatory roles in invertebrates during plastic responses to the environment and (ii) that these effects are contingent upon additional epigenomic and genomic states. This degree of complexity underlying differential GBM’s functions juxtaposes the simple relationship between promoter DM and expression exhibited across vertebrates (Boyes & Bird, 1992). Such complexity is expected, however. In model species for which GBM has been frequently investigated, its potential effects are numerous, interrelated, and remain a point of active debate (Zilberman, 2017). Our results provide additional evidence that GBM has multiple non-mutually exclusive functions in metazoans, extending current knowledge in that GBM’s multivariate effects may be shaped by a multifactorial epigenomic space. While a great deal remains to be uncovered about the gene regulatory roles of GBM in metazoans, several key results arose from our study that may aid in understanding the physiological significance of epigenomic regulation associated with responses to environmental stress and phenotypic plasticity.

The effect of GBM on DE was strongest among genes with high absolute intron DM and low TSS accessibility, with transcript abundance influencing the direction of effect (Fig. 3A). We also observed that introns were significantly more accessible than exons and promoters and that intron methylation shared stronger correlations with the occurrence alternative splicing events than exon methylation (Fig 1B). These feature-dependent effects on both baseline and plastic patterns of gene regulation underscore the unique roles that exon and intron methylation may possess. While our analyses indicate, with strong likelihood, that the regulatory role of intron DM is dependent on TSS accessibility and expression level, the mechanisms by which intron methylation contributes to gene regulation, and how those mechanisms differ from exon DM, remain unresolved.

Previous studies have identified effects of intron DM on DE outside of invertebrates and interactions between GBM and TSS accessibility in their effects on baseline expression, but there is strong potential for such effects to phylogenetically vary. GBM is positively correlated with TSS accessibility measured using a putative approach for several arthropods (Lewis et al., 2020), negatively correlated with TSS accessibility as demonstrated by ATAC-seq in the crustacean *Procrambus virginalis* (Gatzmann et al., 2018) and uncorrelated with TSS accessibility in *S. purpuratus* (Fig. S4). With regard to intron methylation, CpG methylation of large introns can be required for gene expression (Rigal, Kevei, Pelissier, & Mathieu, 2012). Conversely, methylation of first introns can be negatively correlated with gene expression, suggesting a distinct role relative to introns 2 – *n* in plants and vertebrates (Anastasiadi, Esteve-Codina, & Piferrer, 2018; Rose, 2008; Tan, 2010). The mean intron length of *S. purpuratus* is 1.753 kb (Tu, Cameron, Worley, Gibbs, & Davidson, 2012) and, as we have demonstrated, average % CpG methylation is relatively even across introns and exons. Basal invertebrates such as placozoans and sponges exhibit shorter intron lengths than higher order invertebrates such as deuterostomes (McCoy & Fire, 2020). Furthermore, the evenness of CpG methylation between introns and exons of *S. purpuratus* is distinct from most other invertebrates for which exon methylation is greater (Downey-Wall et al., 2020; Lewis et al., 2020; Li et al., 2018) but comparable to other echinoderms (Yang, Zheng, Sun, & Chen, 2020). Despite our support in *S. purpuratus* for the hypothesis that DNA methylation’s gene regulatory roles during plasticity are dependent additional epigenomic and genomic states, it is important to underscore that the nature of intron methylation’s effect on DE and its interactions with TSS accessibility may vary across invertebrate phyla.

DE induced by maternal stress was most positively affected by intron DM among lowly expressed genes with low TSS accessibility that were enriched for molecular functions including ‘nucleotidyltransferase’ and ‘cytoskeletal motor activity’, molecular functions that were also enriched among genes that were differentially methylated following maternal upwelling. ‘Nucleotidyltransferase activity’ was also enriched among genes that were downregulated in response to maternal upwelling. (Strader et al., 2020). Therefore, it is possible that the DM and DE of some gene families were attributed to interactions between methylation, chromatin accessibility, and expression level.

Importantly, intron DM exhibited a negative correlation with DE among genes with low TSS accessibility, but high expression. Bidirectional effects of differential GBM on DE could help explain why past associations between differential GBM and expression have been negligible in invertebrates. Multiple hypotheses may explain how the directionality of intron DM’s association with DE changed according to expression level. Introns possess both enhancing and silencing effects on gene expression across eukaryota (Rose, 2008; Rose, 2018). Intragenic enhancers are predominantly located in introns and their methylation can be negatively correlated with expression in some eukaryotic lineages (Blattler et al., 2014). Enhancer profiling in developing *S. purpuratus* has revealed correlations between enhancer activity and gene expression, indicating that a proportion of highly expressed genes in early prism-stage larvae may correspond to a greater number of activated enhancers (Khor, Guerrero-Santoro, Douglas, & Ettensohn, 2021). Long introns are also enriched with conserved regulatory elements such as transcriptional enhancers (Haddrill, Charlesworth, Halligan, & Andolfatto, 2005; Park et al., 2011), potentially explaining the silencing effect of intron DM we observed among *S. purpuratus* genes with long intron lengths (Fig. S3). Future work should test whether intron hypermethylation induces downregulation of transcripts enriched with intragenic enhancers and whether positive associations between intron DM and DE are attributable to MBD-binding at methylated gene bodies, promoting histone H3K27 acetylation and gene expression (Xu et al., 2021).

### 4.3. Effects of differential methylation on splicing are conditional upon chromatin and genic architecture states

Alternative splicing diversifies the proteome and is an essential and conserved gene regulatory mechanism (Keren, Lev-Maor, & Ast, 2010). Changes in exon use under stress can be attributable to alterations in both constitutive and alternative splicing events (Biamonti & Caceres, 2009). In response to maternal upwelling, differential GBM in *S. purpuratus* larvae interacted with chromatin state and genic architecture to potentially influence alternative splicing and/or exon skipping. A three-way interaction between exon DM, exon accessibility, and genic intron length affected DEU such that (i) the absolute effect of exon DM was strongest among genes with greater total intron lengths and (ii) DEU of poorly accessible exons positively correlated with exon DM while the DEU of accessible exons negatively correlated with exon DM. To our knowledge, our results mark the first evidence in an invertebrate of a significant association between DM and exon inclusion responding to environmental variation.

The effect of total intron length on DEU aligns with observations that alternative splicing is more pervasive among long genes (Flores et al., 2012; Grishkevich & Yanai, 2014), whose size is largely attributable to intron length in *S. purpuratus* (Tu et al., 2012). Long genes involved in cellular structure and cell adhesion are targets of alternative splicing, producing a diversity of protein isoforms that can modify protein-protein complexes that construct or regulate the cytoskeleton, organelle organization, and the extracellular matrix across cell types, developmental stages, and physiological states as evidenced in multiple model systems (Belkin et al., 1997; Exposito, D’Alessio, & Ramirez, 1992; Kalsotra & Cooper, 2011; Leung, Zheng, Prater, & Liem, 2001; O’Leary, Lasda, & Bayer, 2006). Long genes with poorly accessible exons in *S. purpuratus* were enriched with ‘protein-protein dimerization’ MF GO terms and BP terms such as ‘cellular component assembly’ and ‘organelle organization’. Similar to genes whose DE was most affected by DM, these genes were also enriched with signaling receptor and signal transduction functions. ∼65% of exons with the strongest positive associations between DM and DEU (e.g., high genic intron length and low exon accessibility) came from genes involved in organelle organization, cytoskeletal structure, and the extracellular matrix.

The positive correlation between exon DM and DEU in genes with inaccessible exons was expected as hypermethylation at alternatively spliced exons has generally been associated with their inclusion in transcripts (Flores et al., 2012; Shayevitch, Askayo, Keydar, & Ast, 2018). As stated however, the direction of exon methylation’s effect on splicing was influenced by chromatin accessibility; accessible exons exhibited a negative correlation between DEU and exon DM. While our study cannot derive the exact mechanism by which exon DM and accessibility interacted to affect exon use, several lines of evidence demonstrate their joint influence on alternative splicing. For example, the methyl-binding protein MeCP2 links splicing, DNA methylation, and chromatin state. MeCP2 aids in the recognition of alternatively included, methylated exons resulting in a positive correlation between methylation and inclusion among MeCP2-regulated exons (Lev Maor, Yearim, & Ast, 2015). MeCP2 also interacts with nucleosomes, and its genomic positioning is associated with H3K27Me3, a histone mark related to chromatin inaccessibility (Yin et al., 2021). Thus, the negative correlation between exon DM and DEU among genes with inaccessible exons could result from MeCP2 regulation. Methylated exons that are regulated by the transcriptional repressor CTCF experience skipping rather than inclusion. CTCF induces exon inclusion, but its binding to exons is inhibited by DNA methylation, resulting in a negative correlation between methylation and inclusion (Lev Maor et al., 2015). CTCF binding motifs are associated with increased chromatin accessibility (Jain, Ba, Zhang, Dai, & Alt, 2018). Thus, genes with accessible exons for which a positive correlation exists between exon DM and DEU may be regulated by CTCF. CTCF is conserved and expressed in *S. purpuratus* (Gomez-Marin et al., 2015) and MeCP2 is present in the *S. purpuratus* genome assembly and transcriptome.

## 5. Conclusion

Variation in DNA methylation appears to be a component of molecular responses by many invertebrates to predicted global change including ecologically critical, threatened groups such as stony corals (Putnam et al., 2016) and pteropods (Bogan, Johnson, & Hofmann, 2020) or detrimental invasive species (Hawes et al., 2018). Given the heritability of DNA methylation in some invertebrate clades (Liew et al., 2020), exacting its transcriptional and phenotypic consequences is critical for understanding the mechanistic basis of TGP. Our findings (i) provide quantitative support for the hypothesis that gene regulation by differential GBM in *S. purpuratus* is affected by additional epigenomic and genomic states and (ii) indicate that these effects influence both gene expression and mRNA splicing. The majority of ecological epigenomic studies in metazoans have focused on the singular effects of DM on expression, likely due to the predictive power of promoter methylation in vertebrates (Boyes & Bird, 1992). However, DNA methylation is not a silver bullet to predict transcriptional changes by *S. purpuratus* in response to environmental variation. Rather, it is likely one cog in the epigenomic machinery contributing to plasticity in gene expression and alternative splicing. A shift toward integrated studies combining DNA methylation, chromatin accessibility, and genomic/genic architecture may be necessary to accurately quantify non-genetic sources of transcriptional and phenotypic variation in invertebrates and other eukaryotes.

## Supporting information

Supplemental Results

Datasheet 1

Datasheet 2

Datasheet 3

Datasheet 4

## Acknowledgements

We thank Juliet Wong, Logan Kozal, Terence Leach, Jannine Chamorro, and Maddie Housh for their assistance in executing the culturing experiment from which this study is derived. We are also grateful for insight and comments provided by Steven Roberts and his lab on this project’s early results. Lastly, this work would not have been possible without boating and collection assistance provided by Clint Nelson of the Santa Barbara Channel Long Term Ecological Research program and Christophe Pierre, Director of Marine Operations at UC Santa Barbara.

## Author Contributions

GEH conceived the original design and scope of the experiment. SNB, MES, and GEH conceived hypotheses and analytical approaches. MES led experimental execution and prepared DNA and RNA for sequencing. SNB and MES performed bioinformatic analyses. SNB wrote the manuscript with contributions from MES and input from GEH. All authors have read and approved the final manuscript.

## Data Availability

Raw RNA-seq and RRBS fastq files associated with this study are available through the NCBI Short Read Archive under the accession PRJNA548926. Scripts associated with trimming, mapping, and counting RNA-seq reads and CpG methylation are available in the following GitHub repository: https://github.com/mariestrader/S.purp_RRBS_RNAseq_2019. Code corresponding to all analyses reported in this study, their intermediate files, and outputs can be found in the following GitHub repository: https://github.com/snbogan/Sp_RRBS_ATAC.

## Supplemental Material

Supplemental Results – PDF of supplemental figures and tables

Datasheet 1 – Test statistics for differential expression and exon use

Datasheet 2 – Test statistics for differential methylation across CpGs and genomic features

Datasheet 3 – Parameters of exon use ∼ exon number regressions evaluating spurious transcription and alternative TSS

Datasheet 4 – Enriched GO terms according to Mann Whitney U or Fisher’s exact tests

