## Supplemental Results for "Gene regulation by DNA methylation is contingent on chromatin accessibility during transgenerational plasticity in the purple sea urchin"

**Table of Contents:**

A. Supplemental figures referenced in main text (p. 3 – 6)

B. Specification and diagnostics for selected model of differential expression under maternal upwelling as a function of intron differential methylation (p. 7 – 13)

C. Specification and diagnostics for selected model of differential exon use under maternal upwelling as a function of differential exon methylation (p. 14 – 20)

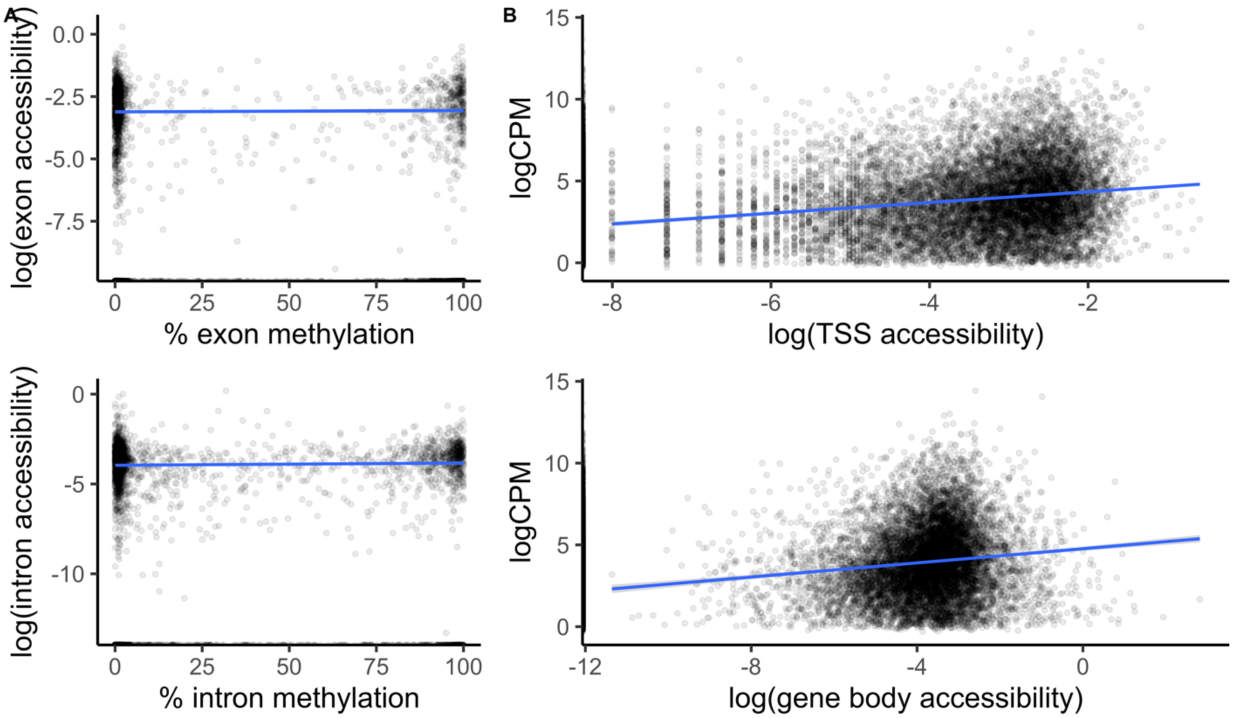


**Figure S1**: Relationships between baseline gene body methylation, chromatin accessibility, and gene expression. (A) % CpG methylation and log chromatin accessibility of exons (top) and introns (bottom) are plotted against one another. (B) TSS accessibility (top) and gene body accessibility (bottom) are plotted against transcript abundance measured as logCPM.

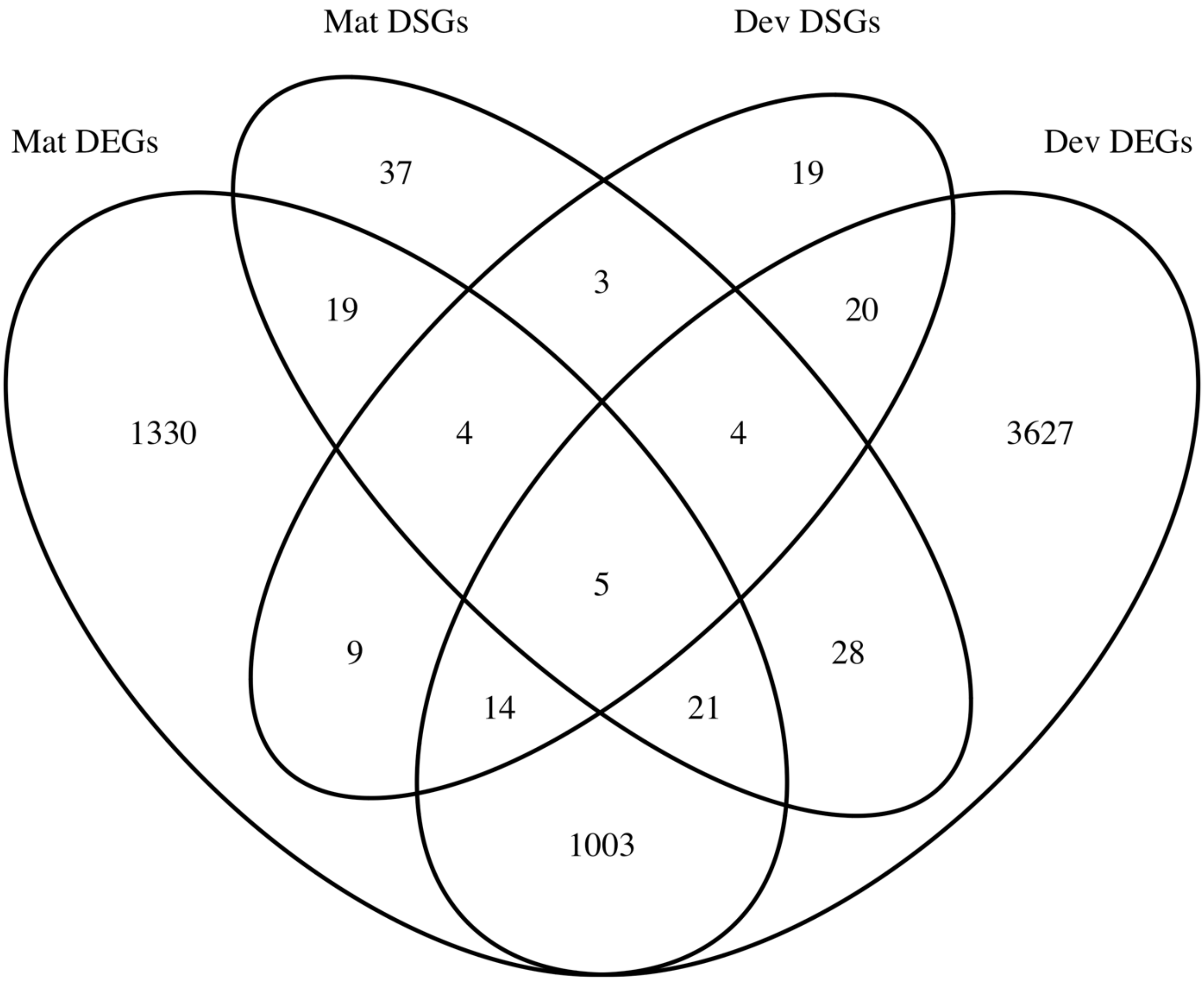


**Figure S2:** Overlap among differentially expressed (DEGs) and differentially spliced genes (DSGs) under maternal and developmental exposure to upwelling (FDR < 0.05).

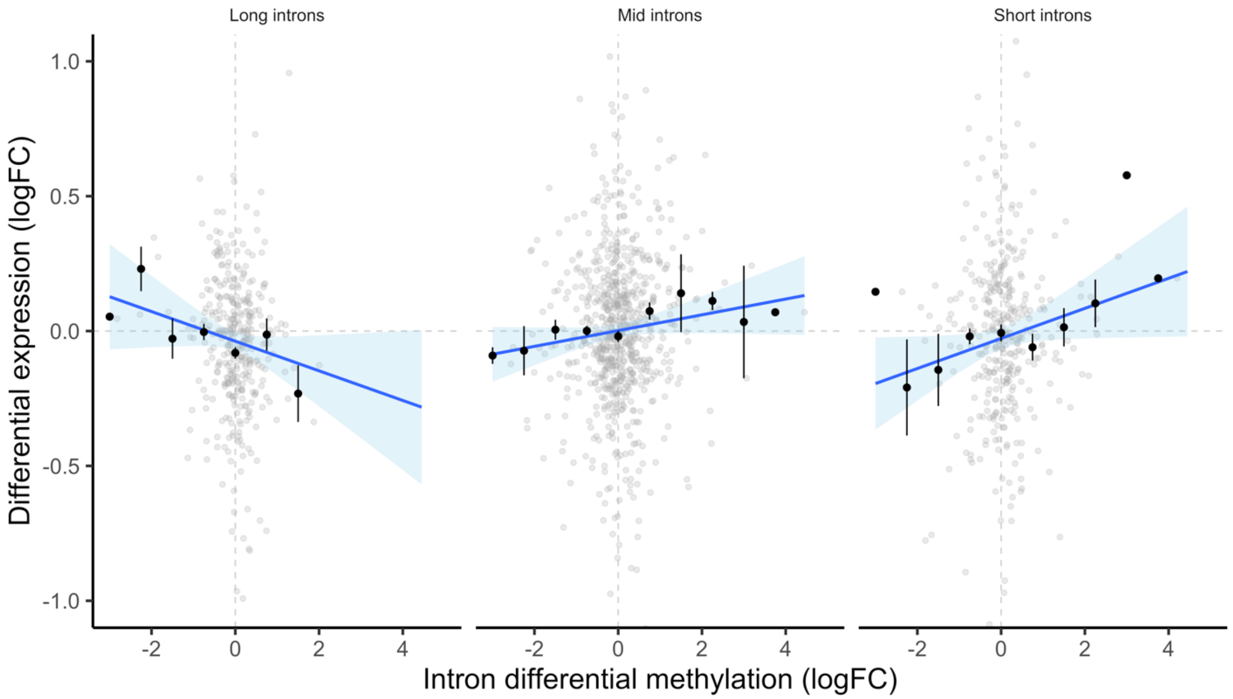


**Figure S3:** Differential intron methylation affected expression conditional upon intron length. Differential gene expression under maternal upwelling is plotted against mean intron differential methylation. Data are grouped based on genic intron length quartiles. ‘Long introns’ and ‘short introns’ denote highest and lowest quantiles. Linear regressions are fitted across observed values. Average logFC across binned intron differential methylation is plotted as black points ± SE.

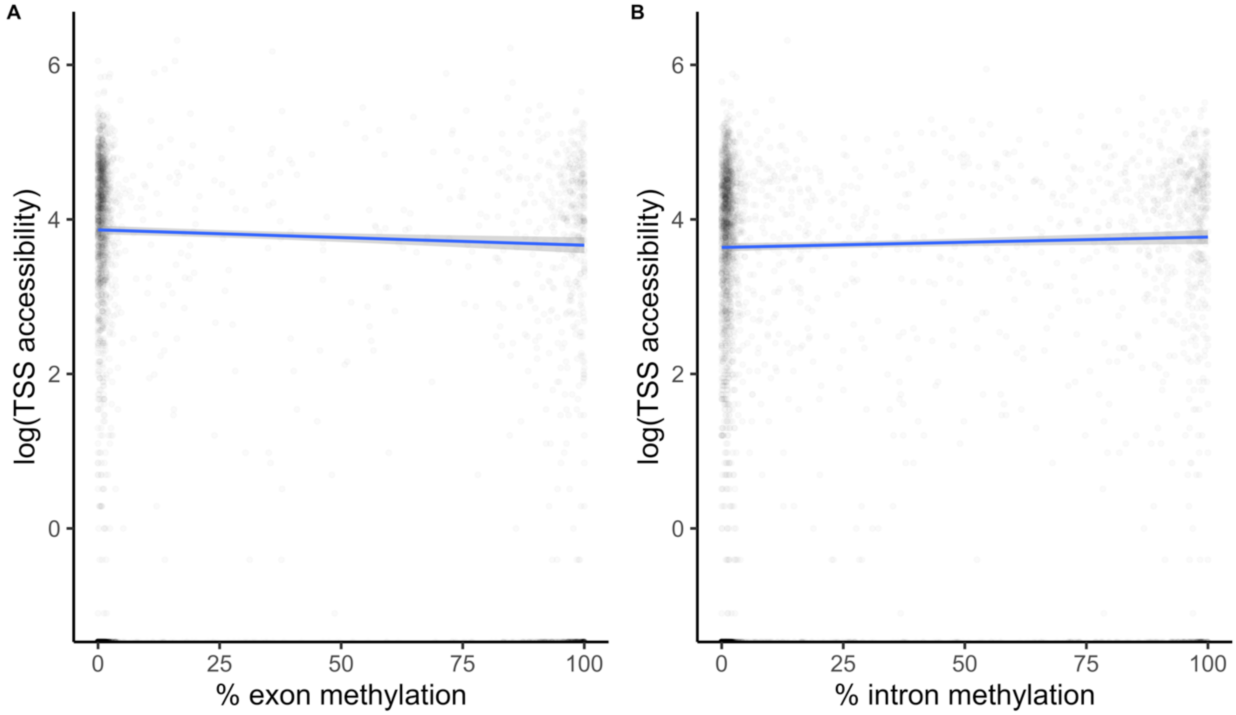


**Figure S4:** Relationships between $\log_{10}$ TSS accessibility and gene body methylation at (a) exons and (b) introns.

### Specification and diagnostics for selected model of differential expression under maternal upwelling as a function of intron differential methylation

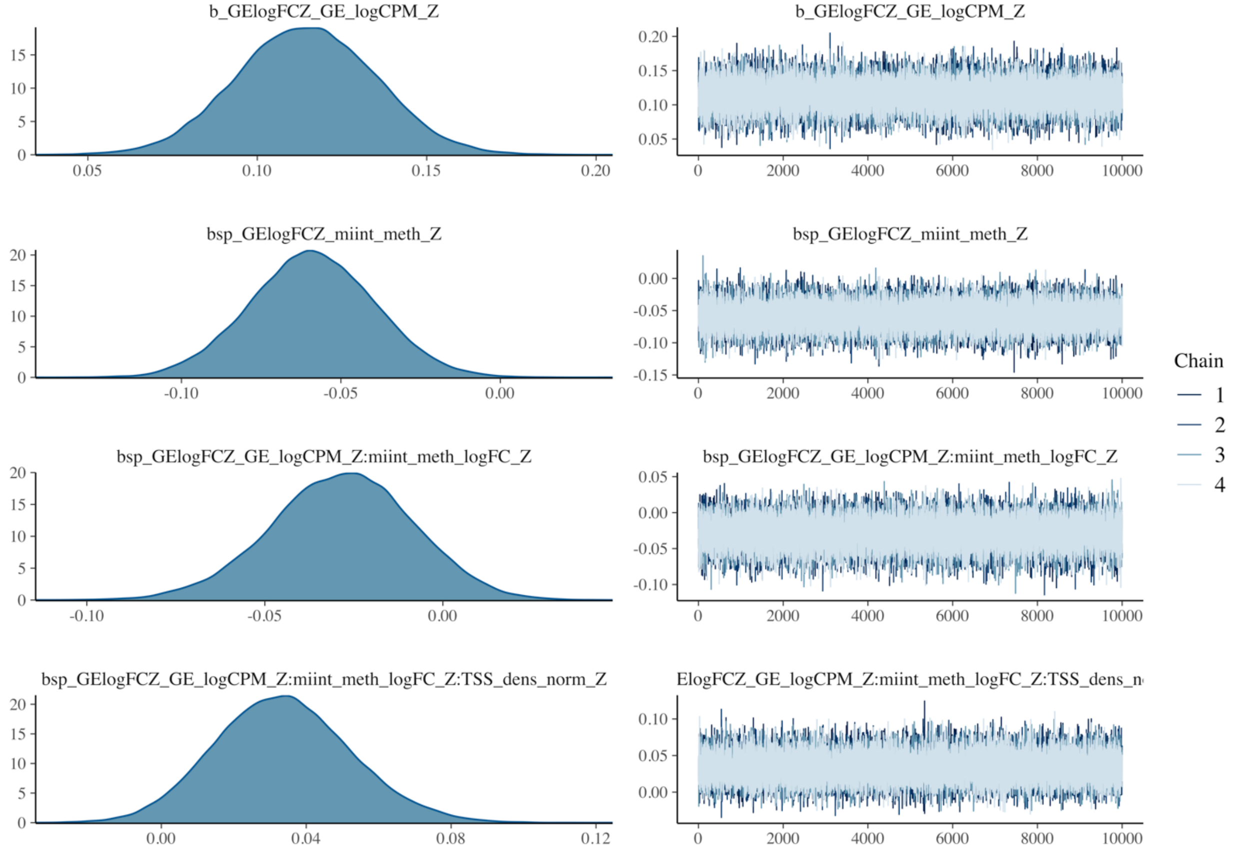


**Figure S5:** Posterior distributions and MCMC chains for β parameters of the selected model predicting differential expression under maternal upwelling as a function of differential intron methylation. "_Z" is appended to the end of parameters that were scaled to Z-scores during model fitting in order to improve run time and convergence of MCMC chains. 'GE_logCPM' denotes logCPM of gene expression. 'int_meth' denotes baseline % methylation of intronic CpGs. 'int_meth_logFC' represents $log_2$FC of differential intron methylation across genes. 'TSS_dens_norm' represents the density of chromatin accessibility at ± 500 bp TSS regions.

### Specification and diagnostics for selected model of differential expression under maternal upwelling as a function of intron differential methylation

| Parameter | 5% interval | 95% interval |
|---|---|---|
| b_Intercept | -4.188403e-02 | 2.428285e-02 |
| b_GE_logCPM_Z | -8.080685e-02 | 1.478402e-01 |
| b_int_meth_Z* | -9.111463e-02 | -2.659821e-02 |
| b_GE_logCPM_Z:int_meth_logFC_Z* | -6.181759e-02 | 3.732449e-03 |
| b_GE_logCPM_Z:int_meth_logFC_Z:TSS_dens_norm_Z* | 3.671764e-03 | 6.537291e-02 |
| sigma | 5.643554e-01 | 6.325130e-01 |
| nu | 2.541751 | 3.388320 |
| Intercept | -4.188403e-02 | 2.428285e-02 |

**Table S1**: Posterior intervals representing probability of direction tests applied to selected model of differential expression under maternal upwelling. Significant effects have posterior probabilities for which >95% of the distribution falls above or below 0. "_Z" is appended to the end of parameters that were scaled to Z-scores during model fitting in order to improve run time and convergence of MCMC chains. 'GE_logCPM' denotes logCPM of gene expression. 'int_meth' denotes baseline % methylation of intronic CpGs. 'int_meth_logFC' represents $log_2$FC of differential intron methylation across genes. 'TSS_dens_norm' depicts the density of chromatin accessibility at ± 500 bp TSS regions. An asterisk denotes methylation parameters modeled with an error term equaling inverse gene-level intron CpG coverage per gene.

**Specification and diagnostics for selected model of differential expression under maternal upwelling as a function of intron differential methylation**

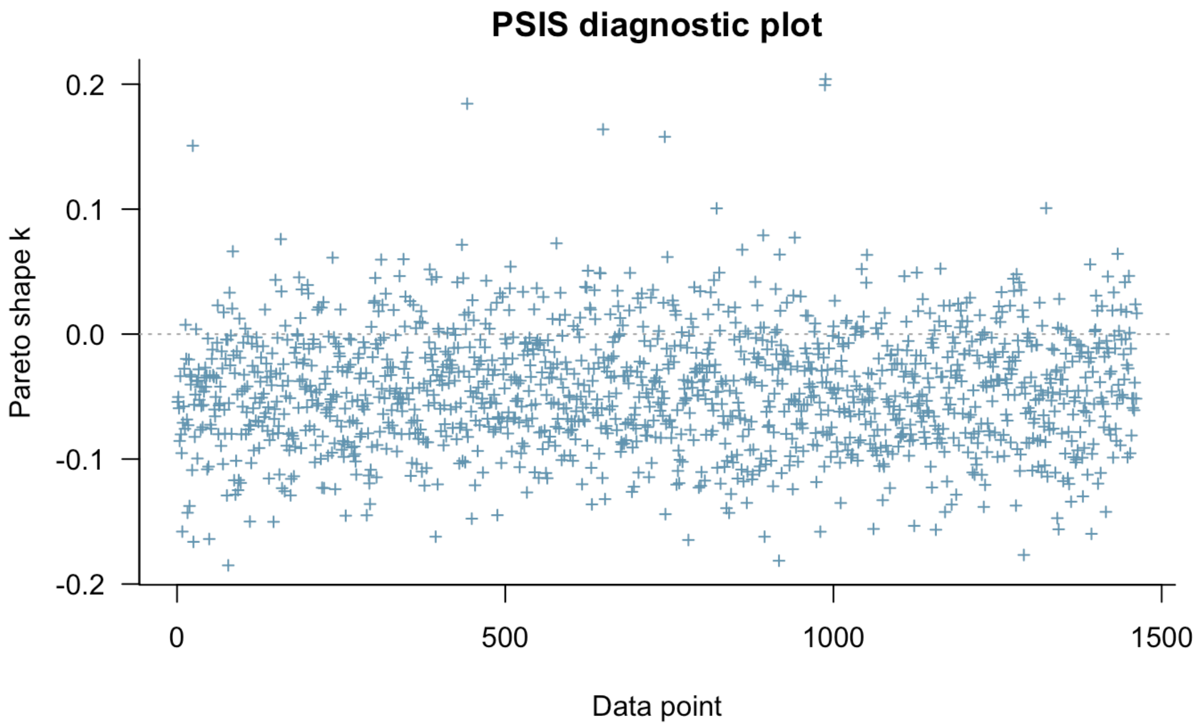


**Figure S6:** Leave-one-out estimates of leverage for observed data fit to selected model of differential expression under maternal upwelling as a function of differential intron methylation. Observations with pareto shape k > 0.4 are deemed to have moderate leverage capable of biasing model fitting. Observations with pareto shape k > 0.7 possess high leverage.

### Specification and diagnostics for selected model of differential expression under maternal upwelling as a function of intron differential methylation

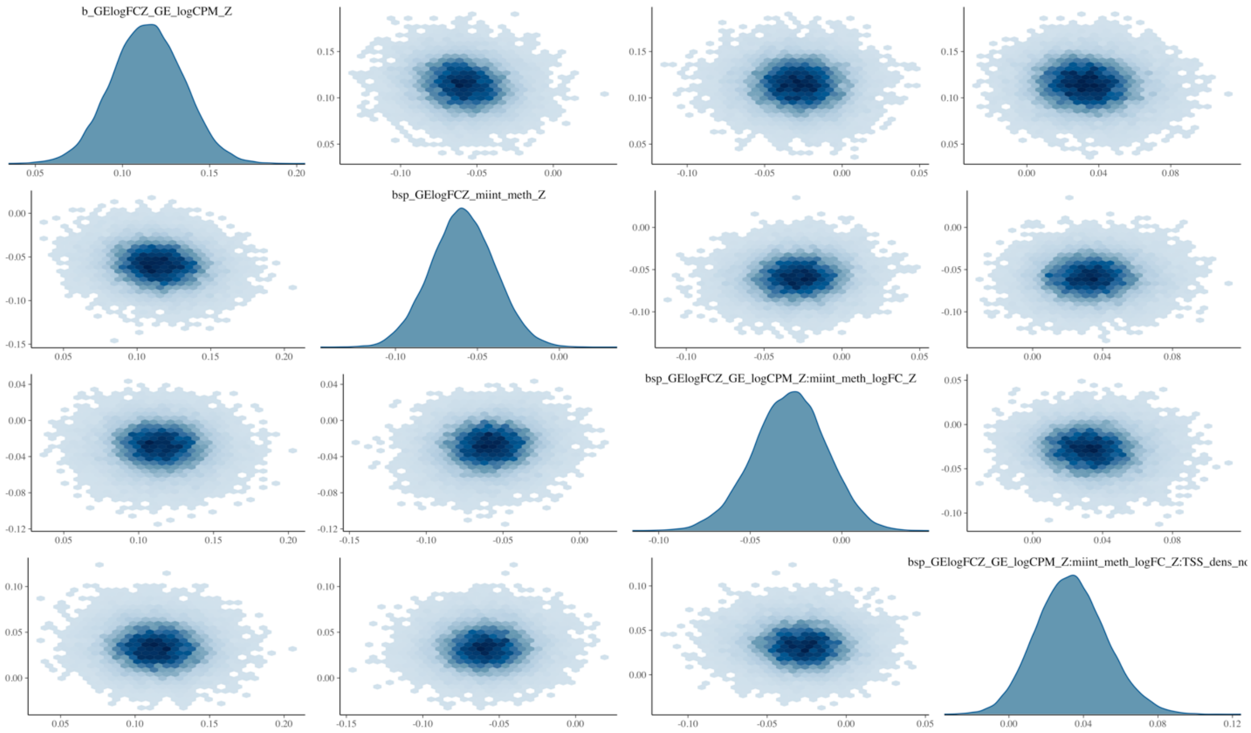


**Figure S7:** Correlation matrix of β posterior draws for fixed effects in selected model of differential expression under maternal upwelling as a function of differential intron methylation. Darker blue depicts greater point density. "_Z" is appended to the end of parameters that were scaled to Z-scores during model fitting. 'GE_logCPM' denotes logCPM of gene expression. 'int_meth' denotes baseline % methylation of intronic CpGs. 'int_meth_logFC' represents $log_2$FC of differential intron methylation across genes. 'TSS_dens_norm' depicts the density of chromatin accessibility at ± 500 bp TSS regions.

### Specification and diagnostics for selected model of differential expression under maternal upwelling as a function of intron differential methylation

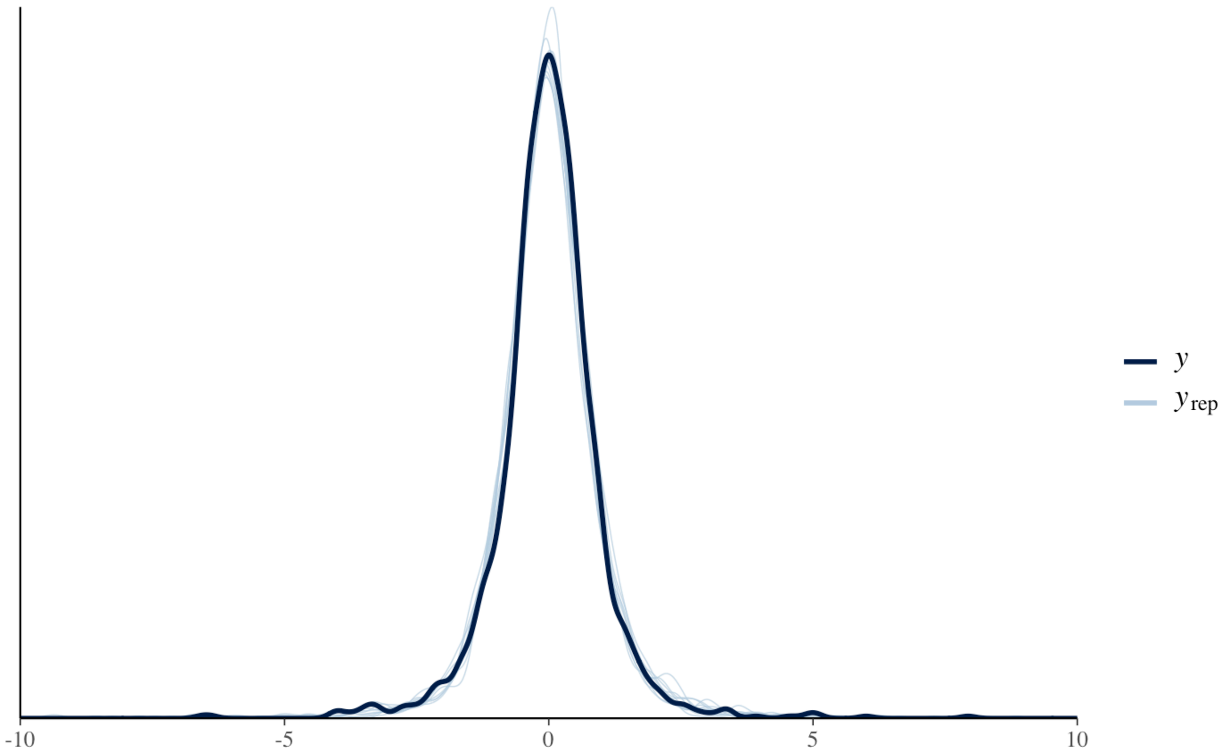


**Figure S8:** Posterior predictive check of selected model predicting differential expression under maternal upwelling as a function of differential intron methylation. The x-axis depicts Z score-scaled differential expression logFC values. The y axis the density distribution of observed and predicted logFC. The black line (γ) depicts the distribution of observed data. Blue lines ($\gamma_{rep}$) depict iterative distributions of model predictions.

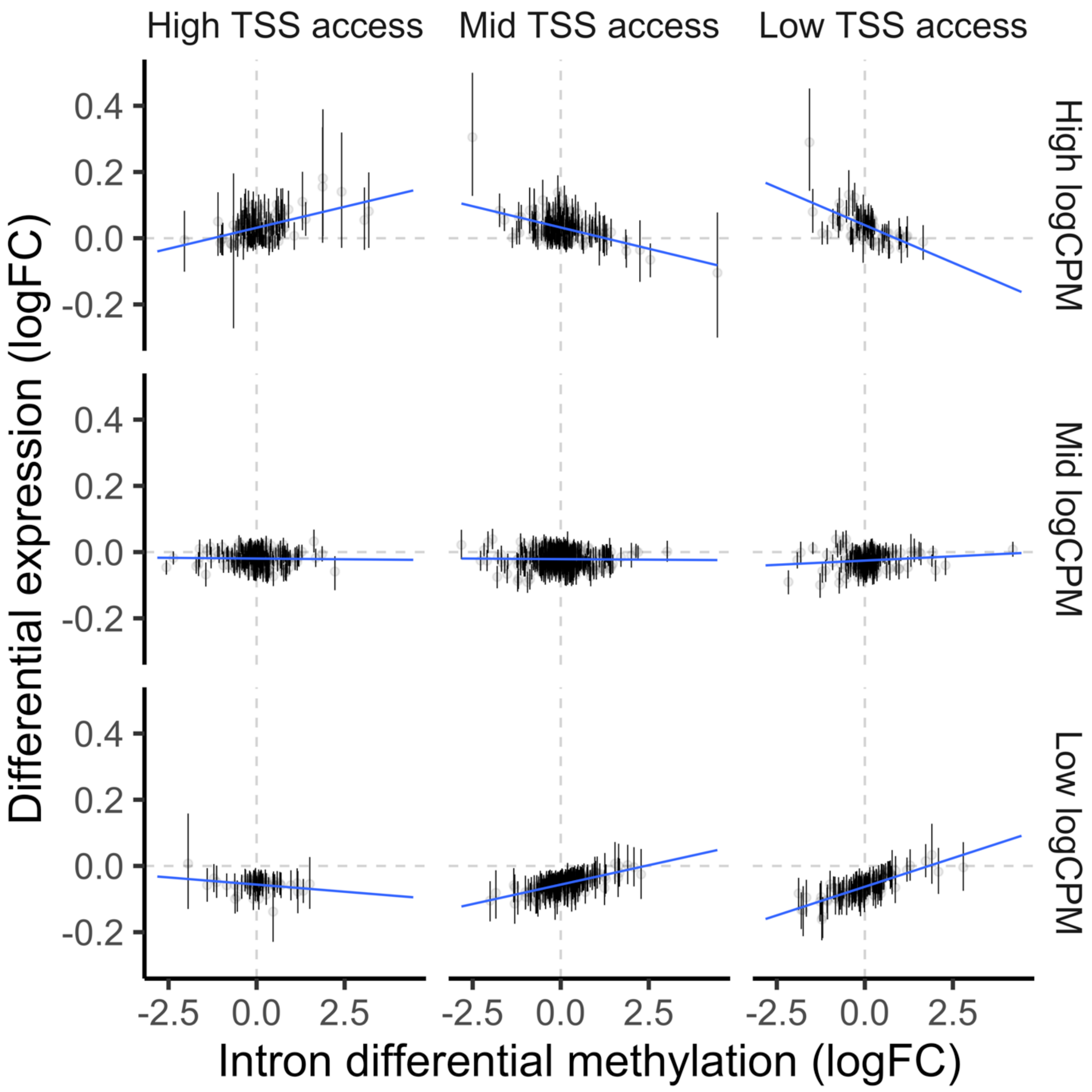


**Figure S9:** Predictions of differential expression under maternal upwelling by selected model relative to intron differential methylation, TSS accessibility (columns), and gene expression level (rows). Individual points represent fitted values ± 95% credibility intervals. 'Low' and 'high' groupings of TSS accessibility and logCPM represent observations in the bottom and top quartiles of these variables.

**Specification and diagnostics for selected model of differential expression under maternal upwelling as a function of intron differential methylation**

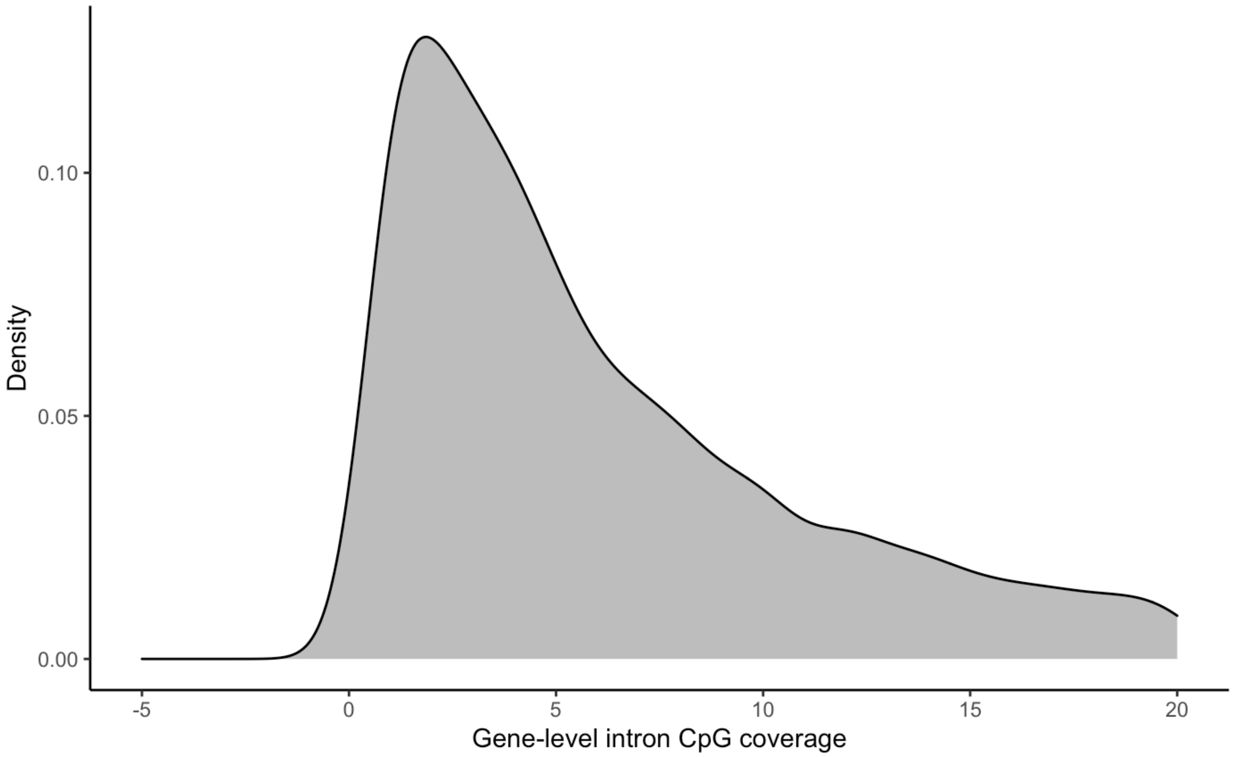


**Figure S10: Gene-level** RRBS CpG coverage of introns post-read count filtering. Mean coverage equaled 14.42 CpGs. Median coverage equaled 6 CpGs.

### Specification and diagnostics for selected model for differential exon use under maternal upwelling as a function of differential exon methylation

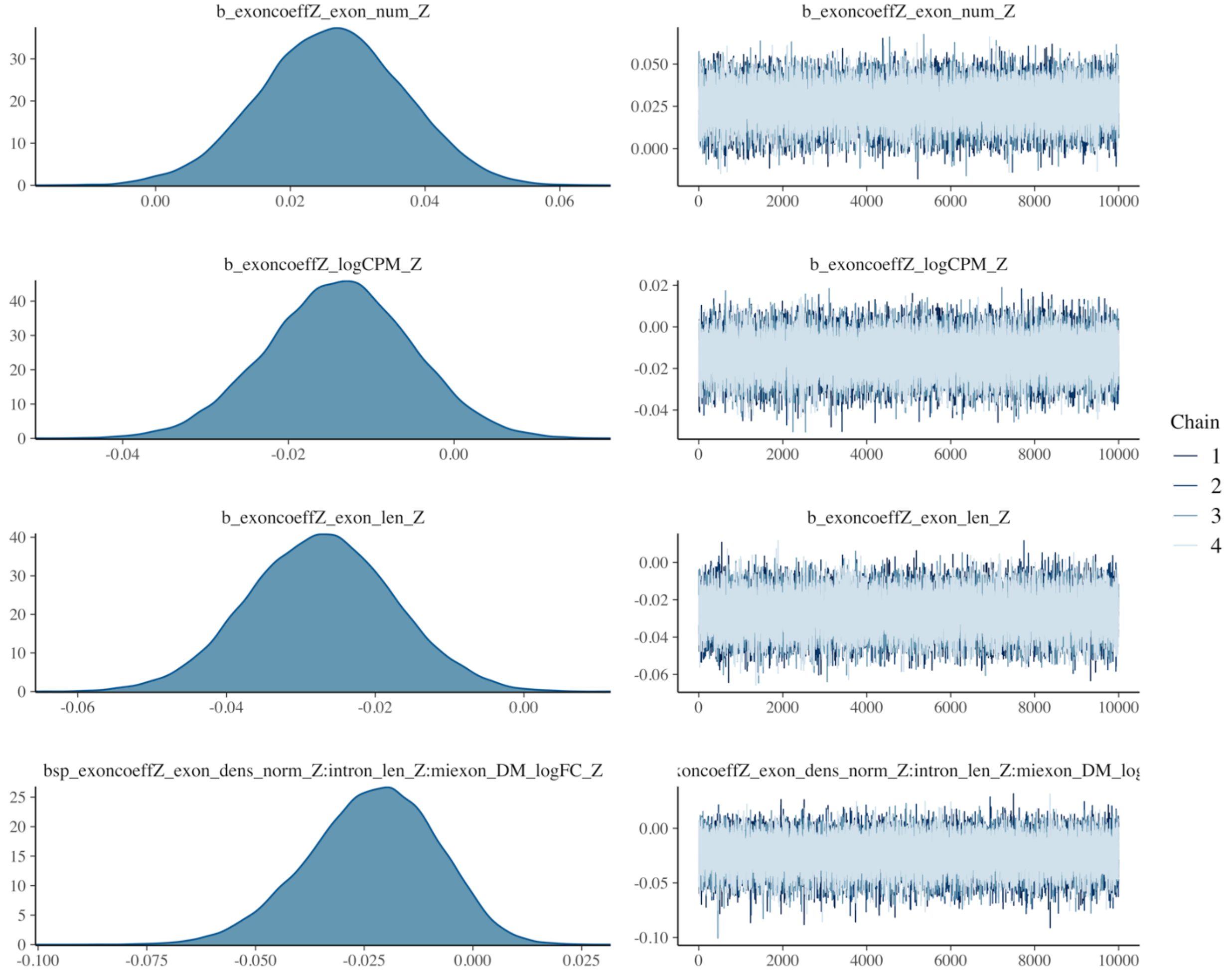


**Figure S11:** Posterior distributions and MCMC chains for β parameters of the selected model predicting differential exon use under maternal upwelling as a function of differential exon methylation. "_Z" is appended to the end of parameters that were scaled to Z-scores during model fitting in order to improve run time and convergence of MCMC chains. 'exon_num' represents exon number. 'logCPM' denotes logCPM of gene expression. 'exon_DM_logFC denotes $log_2$FC values of exon differential methylation. 'intron_len' represents total genic intron length. 'exon_dens_norm' depicts the density of chromatin accessibility at across exons of the associated gene.

**Specification and diagnostics for selected model for differential exon use under maternal upwelling as a function of differential exon methylation**

| Parameter | 5% interval | 95% interval |
|---|---|---|
| b_Intercept | -4.464433e-02 | -1.853375e-02 |
| b_exon_num_Z | 8.625611e-03 | 4.352117e-02 |
| b_logCPM_Z | -2.847342e-02 | 3.777280e-04 |
| b_exon_len_Z _ | -4.325901e-02 | -1.120101e-02 |
| bsp_exoncoeffZ_exon_dens_norm_Z:intron_len_Z: miexon_DM_logFC_Z* | -4.879518e-02 | -2.521101e-04 |
| sigma | 2.415483e-01 | 2.776114e-01 |
| nu | 1.029598 | 1.189859 |
| Intercept | -4.464433e-02 | -1.853375e-02 |

**Table S2**: Posterior intervals representing probability of direction tests applied to selected model of differential exon use under maternal upwelling. Significant effects have posterior probabilities for which >95% of the distribution falls above or below 0. "_Z" is appended to the end of parameters that were scaled to Z-scores during model fitting in order to improve run time and convergence of MCMC chains. 'exon_num' represents exon number. 'logCPM' denotes logCPM of gene expression. 'exon_DM_logFC' represents $log_2$FC of differential methylation at single exons. 'exon_dens_norm' depicts the density of chromatin accessibility at all exons within a gene. An asterisk denotes methylation parameters modeled with an error term equaling inverse exon CpG coverage per gene.

### Specification and diagnostics for selected model for differential exon use under maternal upwelling as a function of differential exon methylation

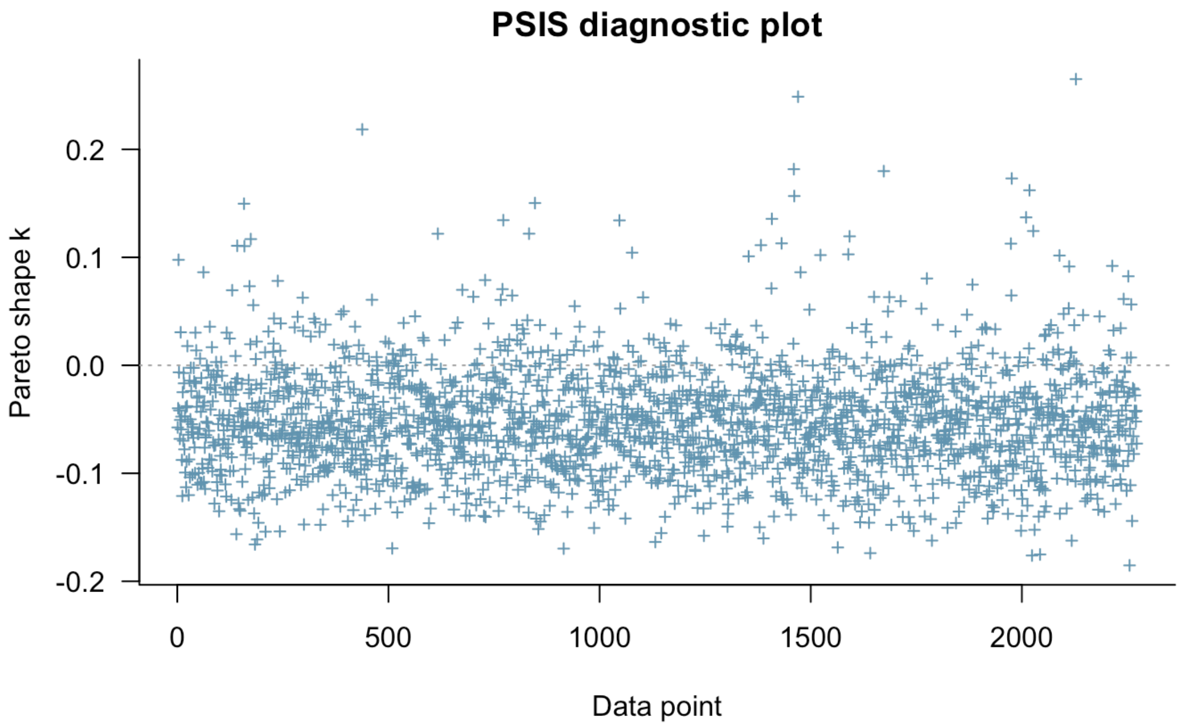


**Figure S12:** Leave-one-out estimates of leverage for observed data fit to selected model of differential exon use under maternal upwelling as a function of differential exon methylation. Observations with pareto shape k > 0.4 are deemed to have moderate leverage capable of biasing model fitting. Observations with pareto shape k > 0.7 possess high leverage.

**Specification and diagnostics for selected model for differential exon use under maternal upwelling as a function of differential exon methylation**

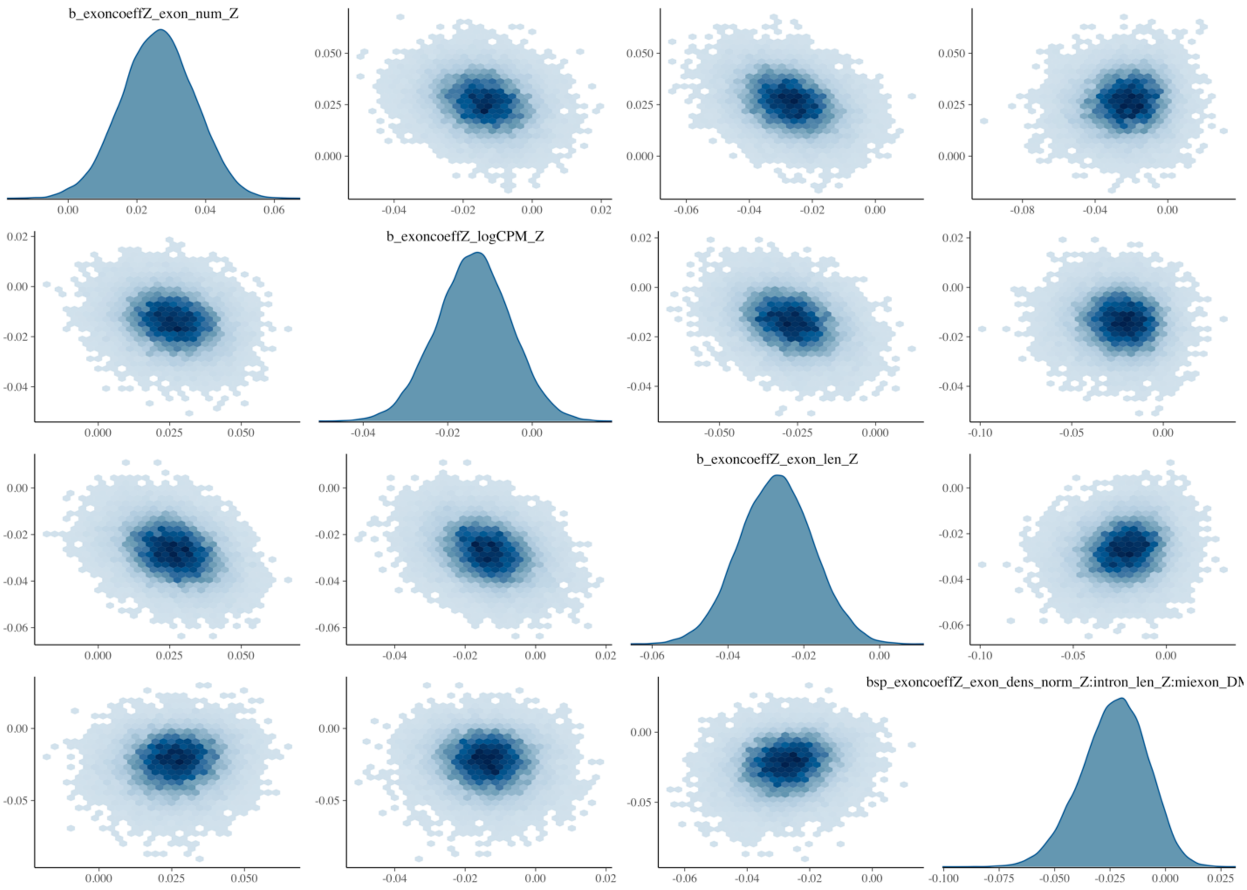


**Figure S13:** Correlation matrix of β posterior draws for fixed effects in selected model of differential exon use under maternal upwelling as a function of differential exon methylation. Darker blue depicts greater point density. "_Z" is appended to the end of parameters scaled to Z-scores during model fitting. 'exon_num' represents exon number. 'logCPM' denotes logCPM of gene expression. 'exon_DM_logFC' represents $\log_2$FC of differential exon methylation across genes. 'exon_dens_norm' depicts the density of chromatin accessibility at exons within a gene. "intron_len" represents total intron length of a gene.

### Specification and diagnostics for selected model for differential exon use under maternal upwelling as a function of differential exon methylation

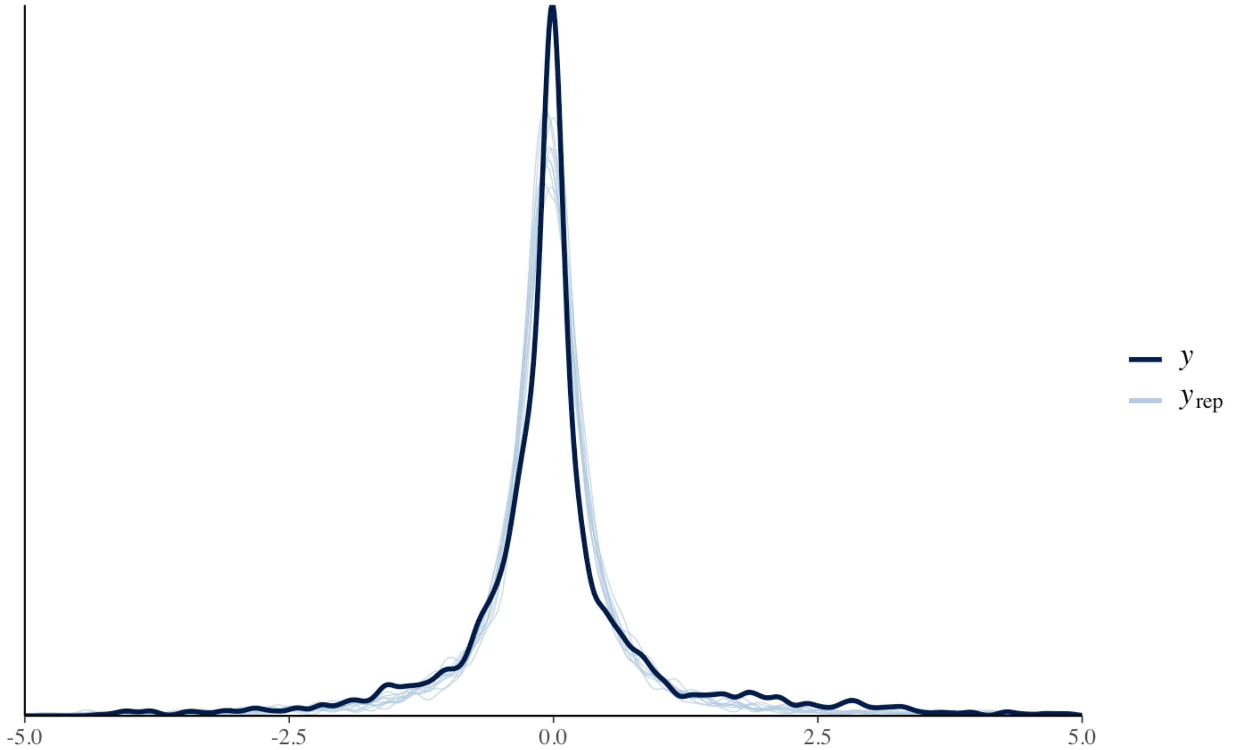


**Figure S14:** Posterior predictive check of selected model predicting differential exon use under maternal upwelling as a function of differential exon methylation. The x-axis depicts Z score-scaled differential exon use ΔlogFC values. The y axis the density distribution of obersved and predicted ΔlogFC. The black line (γ) depicts the distribution of observed data. Blue lines ($\gamma_{rep}$) depict iterative distributions of model predictions.

#### Specification and diagnostics for selected model for differential exon use under maternal upwelling as a function of differential exon methylation

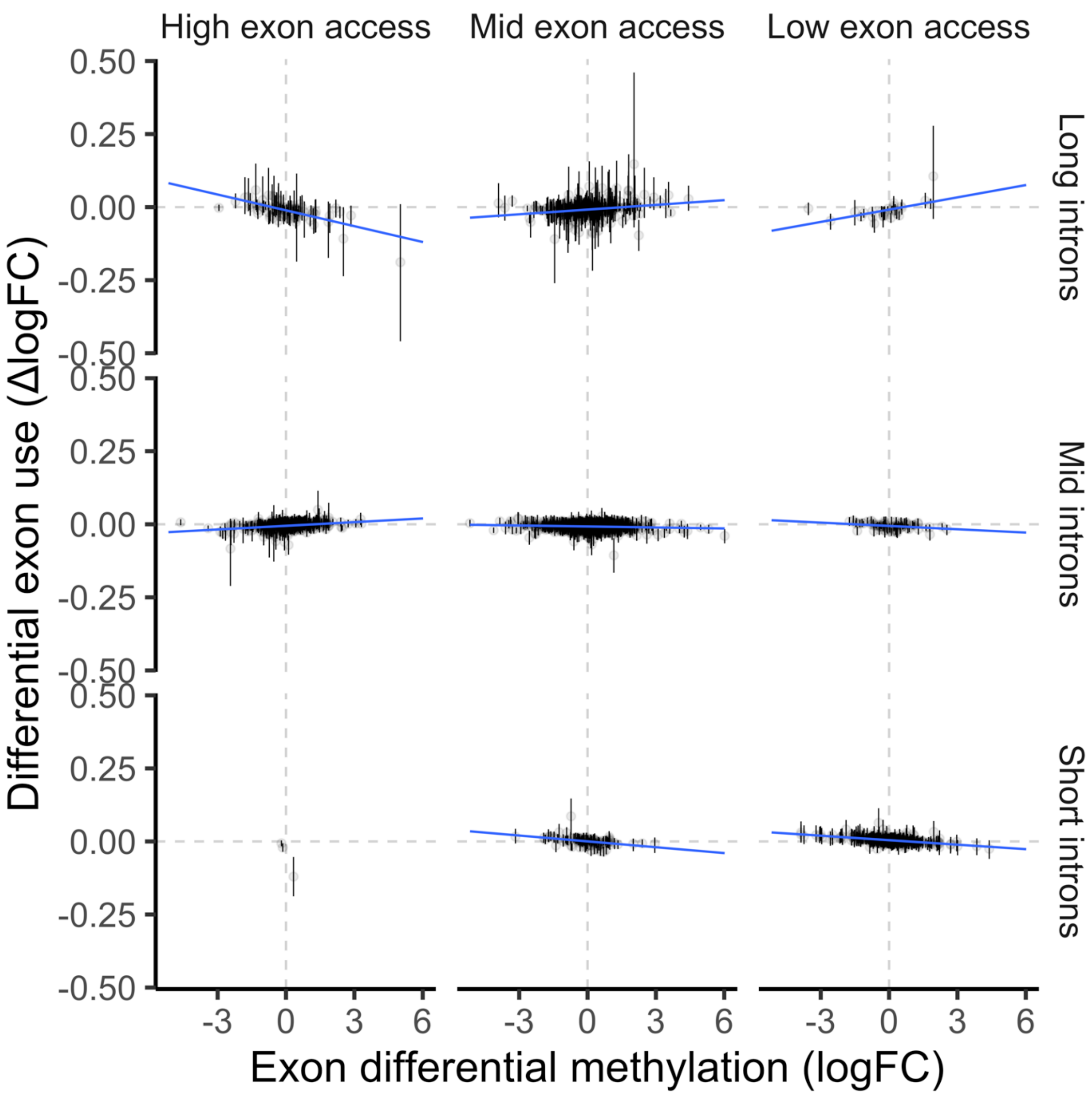


**Figure S15:** Predictions of differential exon use under maternal upwelling by selected model relative to exon differential methylation, exon accessibility (columns), and total genic intron length (rows). Individual points depict fitted values per exon ± 95% credibility intervals. 'Low' and 'high' groupings of exon accessibility and intron length represent observations in the bottom and top quintiles of these variables.

**Specification and diagnostics for selected model for differential exon use under maternal upwelling as a function of differential exon methylation**

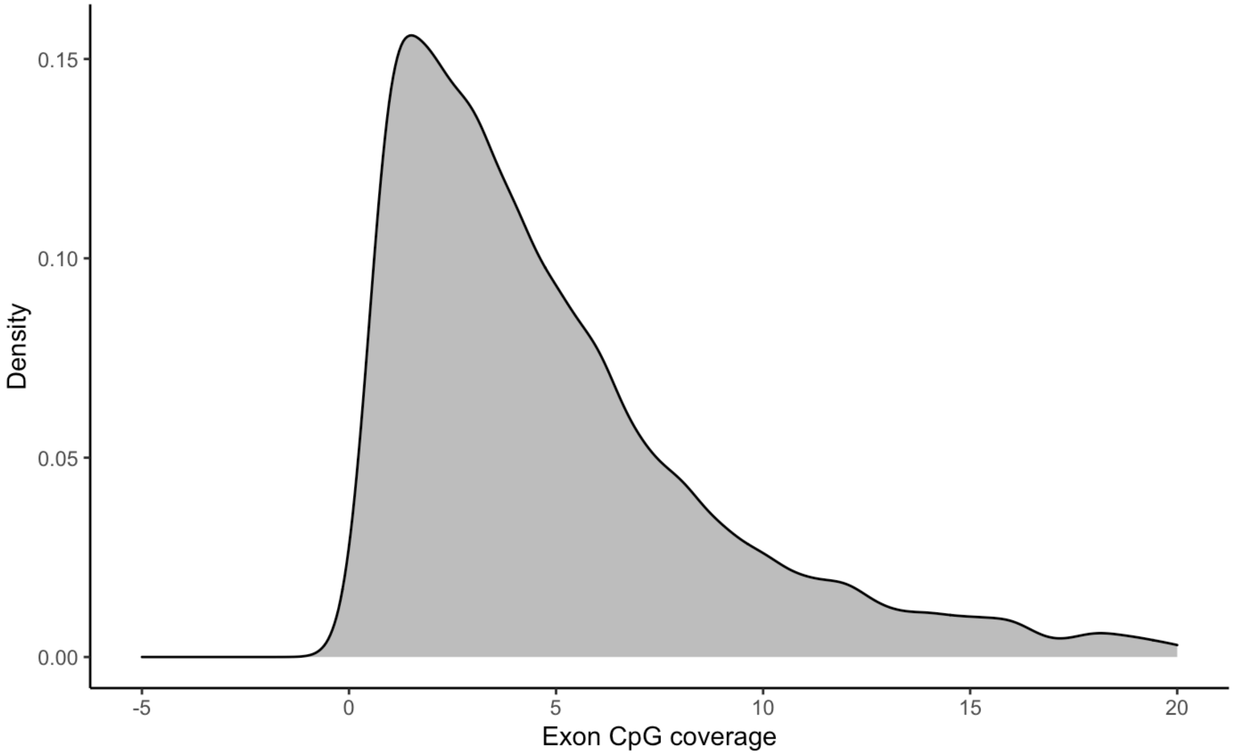


**Figure S16:** RRBS CpG coverage of individual exons post-read count filtering. Mean coverage equaled 5.42 CpGs. Median coverage equaled 4 CpGs.
